# Nanopore-based cell-free DNA fragmentation and methylation profiles from the cerebral spinal fluid of patients with lung cancer brain metastases

**DOI:** 10.1101/2025.07.28.667300

**Authors:** Tianqi Chen, Xiangqi Bai, Georgiana Burnside, Thy Trang Hoang Trinh, Melanie Hayden Gephart, Billy T. Lau, Hanlee P. Ji

## Abstract

**Background:** Non-small cell lung cancer (**NSCLC**) patients with brain metastases (**BMET**) have a poor prognosis. Cerebrospinal fluid (**CSF**) is a source of cell free DNA (**cfDNA**) from the brain and its methylation and fragmentation properties may be an indicator of NSCLC-BMET.

**Methods:** We applied a nanopore single-molecule sequencing approach to characterize the fragmentation, methylation and hydroxymethylation patterns present in CSF-derived cfDNA from NSCLC-BMET patients (N=15). We compared the cancer cfDNA finding to non-cancer healthy controls (N=11) and their CSF cfDNA. We also compared the fragmentation patterns between CSF-derived cfDNA and plasma-derived cfDNA.

**Results:** We observed enriched mono-nucleosome levels and significantly higher mono-/trinucleosome ratios in cancer patients. Comparison with plasma-derived cfDNA further confirmed the unique fragmentation features of CSF-derived cfDNA. Distinct methylation and hydroxymethylation patterns were observed between cancer and control CSF samples. We observed significantly lower degree of hydroxymethylation in cancer patients compared to healthy controls and the affected genes had different pathway profiles.

**Conclusions:** CSF cfDNA in patients with NSCLC-BMET had a distinct profiles of DNA fragmentation, methylation and hydroxymethylation.

## BACKGROUND

Malignancies like lung cancer, breast cancer and melanoma frequently spread to the brain. Lung cancers have the highest risk for brain metastasis (**BMET**) with non-small cell lung cancer (**NSCLC**) accounting for 55% of BMET cases [1]. Approximately 10-20% of NSCLC patients show metastases at diagnosis and over half will develop BMET during their clinical course [2, 3]. NSCLC patients with BMET have a particularly poor prognosis. The median overall survival at 29 months [4]. A common NSCLC genetic feature are somatic *EGFR* mutations which are critical oncogenic drivers in NSCLC [5] and indicative of a specific NSCLC molecular subtype.

*EGFR* mutations are also associated with the presence of multiple metastatic sites, tumor recurrence and a significantly higher risk of BMET [6]. Over 60% of metastatic NSCLC patients with EGFR mutations have BMET.

Tumor cells release their DNA into the blood and other fluids. This analyte is called circulating tumor DNA (**ctDNA**). The ctDNA is a typically a small fraction of the total cell-free DNA (**cfDNA**) circulates in the blood and other body fluids. DNA sequencing of the cfDNA can identify the ctDNA fraction and reveal cancer-specific genomic features such as somatic mutations.

However, the analysis of cfDNA for detecting cancer mutations from the blood has limited sensitivity in identifying cancers located in the brain including BMETs. This may be due to an anatomical structure called the blood-brain barrier that limits the circulation of ctDNA into the bloodstream. Citing an example, for patients with gliomas (a type of brain tumor), less than 10% have detectable ctDNA found in the blood [7]. However, another source of cfDNA is cerebrospinal fluid (**CSF**), a liquid medium which surrounds the brain. The CSF provides buoyancy to neuronal tissue, enables homeostasis to function and facilitates the elimination of waste. Metastatic tumors to the brain release their DNA into the CSF with higher amounts than are found in plasma. Several studies have shown that cancer mutations from brain tumors were detected with greater yield in CSF cfDNA than blood [8, 9], and even the primary tissue [10]. As a result, CSF is a potentially better source for characterizing ctDNA from brain metastasis.

Cell-free DNA has distinctive epigenetic features such as specific patterns of fragmentation and methylation. Both cfDNA and ctDNA fragments are of varying length that directly reflect nucleosomal organization and accessibility to endonucleases. DNA methylation is a chemical modification of an added methyl group in the cytosine as 5-methylcytosine (**5mC**) and frequently subject to tumor-specific differences. The DNA sequences most affected by methylation occur in CpG rich promoter regions, leading to the silencing of gene expression. The 5- hydroxymethylcytosine (**5hmC**), an oxidization product of 5mC, is an intermediate step of DNA demethylation and gene activation. This chemical modification is an epigenetic marker showing reduced levels in many cancers [11]. Moreover, the 5mC and 5hmC patterns are specific to the tissue of origin and cancer types. However, for CSF-derived cfDNA in patients with brain metastases, we know very little about their patterns of DNA fragmentation, 5mC or 5hmC patterns. The cfDNA profile for 5hmC is also challenging to measure given the low levels present in tissues (0.14% in lung, 0.67% in brain) [12].

We applied single-molecule, nanopore sequencing to the CSF cfDNA from patients with metastatic NSCLC. Nanopore sequencing provides multiple DNA features from the same molecule without any intervening molecular processing, PCR amplification or elaborate bioinformatic manipulations [13]. Unlike short read sequencers, there are no restrictions on the length of DNA which can be sequenced. This allows nanopore sequencing to provide longer DNA reads representative of multiple nucleosomes beyond 500 bp read lengths. Also, nanopore sequencing directly identifies cfDNA methylation without chemical or biochemical cytosine conversion. The passage of 5mC and 5hmC DNA through the nanopore generates a unique electrical signal compared to unmodified DNA; a methylated base is detected with machine learning algorithms at high accuracy [14, 15]. This approach does not use PCR amplification. As a result, we avoid GC-biased amplification skews from bisulfite conversion and enabled direct single-molecule counting of cfDNA. The measured methylation profiles directly reflect the native single-molecule state of the cfDNA.

For this study, we conducted a study of CSF-derived cfDNA from patients with NSCLC brain metastases compared to CSF from healthy controls. We determined the CSF-based cfDNA patterns of DNA fragmentation, 5mC methylation and 5hmC hydroxymethylation patterns specific to brain metastasis in patients with NSCLC.

## METHODS

### Samples

We obtained informed consent from all patients based on a protocol approved by Stanford University’s Institutional Review Board (IRB protocol numbers 12625 and 62499). Our cohort consisted of 11 patients with NSCLC brain parenchymal or leptomeningeal metastases and 11 individuals without cancer that provided controls. CSF samples were mostly from lumbar puncture (*N* = 13) procedures, VP shunt (*N* = 1) and Ommaya reservoir (*N* = 1). All samples were stored at − 80 °C before processing. We also used 23 plasma samples obtained from metastatic NSCLC patients (Precision for Medicine). From this set, four patients had NSCLC- BMET while the rest 19 had other metastases from NSCLC. One hundred ninety-eight normal plasma samples were obtained as single aliquots in 1-2 mL cryovials. Thirteen normal plasma samples (Discovery Life Sciences) were used as additional controls. The CSF samples were centrifuged at 1,000 rcf for 10 mins at 4 °C. The supernatant was transferred as 1-2 mL aliquots and stored at -80 °C before sequencing.

### Processing and sequencing CSF cfDNA

Extracted cfDNA was obtained from CSF and plasma samples on KingFisher Apex (Thermo Fisher Scientific) using MagMAX cfDNA isolation kit (Thermo Fisher Scientific) and the standard protocol. For samples with large plasma volume (> 600 µL), the extraction was performed in KingFisher 24 Deep-Well plate (Thermo Fisher Scientific) by a script MagMAX cfDNA-2mL- Flex.bdz (available online) for 2 mL samples and a modified script for 1mL samples. For samples with small volume (< 600 µL), the extraction was performed in KingFisher 96 Deep-Well plate (Thermo Fisher Scientific). Briefly, 1 to 2 mL CSF or plasma was added into one or two 24 DW plates (or 600 µL into two 96 DW plates) as a sample plate with lysis buffer and beads from the kit. The different plates were placed into the KingFisher Apex machine in order, and the script was run. The cfDNA yields were measured by Qubit (Thermo Fisher Scientific). Eluted cfDNA was transferred to Eppendorf DNA LoBind Tubes and stored at -80 °C before library preparation.

We developed an optimized protocol for generating sequencing libraries using plasma-derived cfDNA [13] and was adapted to CSF-derived cfDNA in this work. Briefly, 25-50 μL of extracted cfDNA (corresponding to 0.3-1 ml of CSF) was used as sample input, which underwent end-repair and A-tailing with conditions of 20 °C for 30 min and 65 °C for 30 min (Roche KAPA HyperPrep kit). We ligated native barcodes using 5 μL of each barcoded adapter (SQK- NBD114.24, Oxford Nanopore Technologies) following the standard reaction volumes in the KAPA HyperPrep workflow. We used a thermocycler for the ligation step for 4.5 h incubation at 20 °C before holding at 4 °C overnight to maximize the ligation yield.

After the ligation step, 1 µL EDTA (0.5 M, pH 8.0, Thermo Fisher Scientific) was added into each sample to stop the reactions. Two bead cleanups at 0.8x and 1.5x were performed. First, 88 μl of Mag-Bind Total NGS beads (Omega Bio-Tek) were added and mixed to each reaction. After incubation for 5 min, the beads were magnetized and washed with 80% ethanol using a DynaMag separation rack (Thermo Fisher Scientific) before eluting in 50 μL of 10 mM Tris–HCl pH 8.0 buffer. All samples were pooled together during the elution. Then we performed a second bead cleanup step with 75μL Mag-Bind Total NGS beads (1.5x) and the same magnetic rack procedure. The elution solution was 30 μL 10 mM Tris–HCl pH 8.0 buffer. 10 uL of native adapter was used for the adaptor ligation for 1.5 h at room temperature. Subsequently, we mixed in 150 μl of Mag-Bind Total NGS beads (1.5x) and incubated for 5 min. As in the standard protocol, we washed the beads with 200 μL SFB buffer (Oxford Nanopore Technologies) with gentle tube flicking to resuspend the beads during the wash steps. The beads were resuspended in 33 μL EB buffer (Oxford Nanopore Technologies) and cfDNA eluted. We used 1 μL eluted cfDNA for quantification with a Qubit instrument (Thermo Fisher). The eluted DNA library was stored at 4 °C before sequencing.

We performed sequencing with Oxford Nanopore Technologies’ PromethION 24 instrument using R10.4.1 flow cells. After quantification of the final pooled library, we calculated its molarity. The entire volume of the sequencing library was loaded onto the flow cell.

Sequencing runs had a duration of 72 h.

### Sequence data analysis

The raw pod5 sequencing data were processed using the “super accuracy” basecalling with Dorado v0.5.3 (Oxford Nanopore Technologies, https://github.com/nanoporetech/dorado) with the “dna_r10.4.1_e8.2_400bps_sup@v4.3.0” or “dna_r10.4.1_e8.2_400bps_sup@v4.1.0” model, depending on data sampling frequency of the original run, and demultiplexed for each barcode. The quality score filtering was a value of 7. The GRCh38 reference was used for alignment. The output consists of sequence alignment bam file for each sample containing 5mC and 5hmC information. Bam files were processed by modkit v0.2.6 (Oxford Nanopore Technologies, https://github.com/nanoporetech/modkit) to generate BedMethyl files with 5mC and modified base calls. The calls were extracted into 2 files: 5mC only and 5hmC only. Before further processing, the BedMethyl and bam files were sorted and indexed with samtools [16].

We determined the genome-wide average methylation or hydroxymethylation status of sequenced cfDNA. This involved averaging all 5mC or 5hmC values across all sequenced CpG sites that had at least one read (coverage > 0). To determine nucleosome enrichment, we randomly selected 100,000 reads from each sample’s sequence-aligned bam file, tabulated the estimated fragment size derived from the post-alignment length of the read. The fragment sizes were 167 bp, 330 bp and 500 bp which corresponded to the mono-, di-, and trinucleosome size fragment respectively. To calculate the cutoff values for each type of nucleosomes, we averaged the fragment sizes of mono- and dinucleosomes and di- and trinucleosomes.

Therefore, fragments with size 0-248.5 bp were identified as mono-nucleosome. Fragments with size 248.5-415 bp were identified as dinucleosomes. Fragments with size 415-600 bp were identified as trinucleosomes. Nucleosome ratios of mono-/di-, mono-/tri-, and di-/trinucleosomes were calculated based on the counts of each nucleosomes for each sample. Similar subsampling approach and nucleosome classifications were used in our previous work [13].

We determined CpG-site level methylation for all sequenced cfDNA samples. First, we identified the CpG sites which were covered by at least two cancer samples and two control samples. A *t*-test was performed to compare methylation per CpG site between the non-cancer-derived cfDNA and cancer patient-derived cfDNA. We used FDR-based multiple testing correction to determine statistically significant differences in CpG-site level methylation. We used a cutoff of q < 0.01. Then, these CpG sites were mapped to genes by intersecting their genomic coordinates with GENCODE v38 [17], which includes all coding exons and introns.

Finally, grouping by gene-level annotations, we calculated the average methylation per gene of annotated protein coding genes. For the promoter region analysis, we used two annotations: 1) establishing a window beginning 2 kb upstream and 500 bp downstream of transcription start site (**TSS**) while maintaining strand specificity. The same annotation schema was used in another study [18] and similar ones of -1kb to 500 bp observed in other work [19], and 2) promoter annotation release 112 on GRCh38 by Ensembl defined as 1kb upstream of TSS. The same analysis strategy was applied to 5hmC files to study CpG-site level hydroxymethylation.

### NSCLC methylation patterns

We compared our cfDNA methylation data with the Illumina 450K methylation array data from TCGA_LUAD and statistics about false discovery rates (**FDRs**). The data was available from the CanMethdb database [20]. A threshold of FDR-corrected q ≤ 0.05 was used to identify significant genes. We downloaded the identified CpG-target gene pairs, lifted over to the reference genome GRCh38 and overlapped this information with specific genes of interest from our analysis. We also compared our cfDNA methylation data with the data of open chromatin sites in the promoter region from TCGA_LUAD by ATAC-Seq [21]. This public data consisted of 22 tumor samples, each from a different patient. A threshold of FDR-corrected q < 0.01 was used to identify genomic regions of interest. The promoter annotation was defined as 1 kb upstream and 100 bp downstream of TSS. The GRCh38 was used as the reference genome. We overlapped the promoter genes in the public data with the ones that were significant in our CSF cfDNA results.

## RESULTS

For this study we developed an approach with nanopore single molecule DNA sequencing. It can be applied to cfDNA from any source such as CSF. We used a highly efficient ligation process for a genome-wide sampling [13]. The advantages of this method include: (1) direct sequencing of the cfDNA contents without any PCR amplification skewing or errors; (2) directly methylation calls from the cfDNA that avoids bisulfite chemical or biochemical conversion processing; (3) fragmentation patterns from cfDNA that identifies nucleosome organization without the use of bioinformatic assembly used in short reads.

An overview of the study is shown in **Figure 1**. Description of the clinical cohort is provided in **Table 1**. We evaluated the CSF of 11 patients with metastatic NSCLC, either involving the brain parenchymal, tumors growing in the brain tissue or leptomeningeal metastasis which involves layers of tissue surrounding the brain and spinal cord, namely the arachnoid mater and pia mater. Among the patients with NSCLC BMETs, seven patients had NSCLC *EGFR* mutations and four were *EGFR* wildtype with other type of driver mutations (**Table 1**). In addition, we had CSF from 11 individuals without cancer.

**Figure 1.**
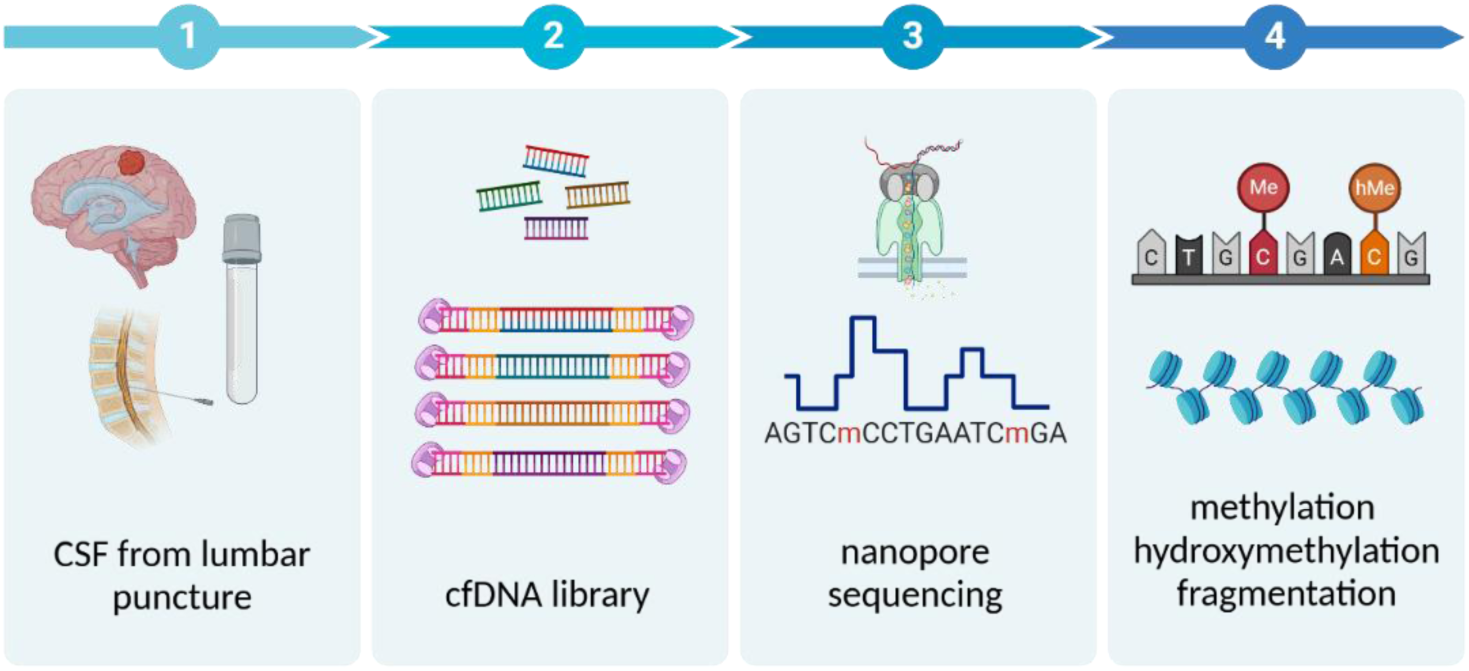
Schema for analyzing CSF cfDNA with nanopore sequencing. The cfDNA was extracted from CSF samples followed by library preparation. Nanopore sequencing was performed. Profiles of cfDNA fragmentation, 5mC and 5hmC were analyzed.

**Table 1.**
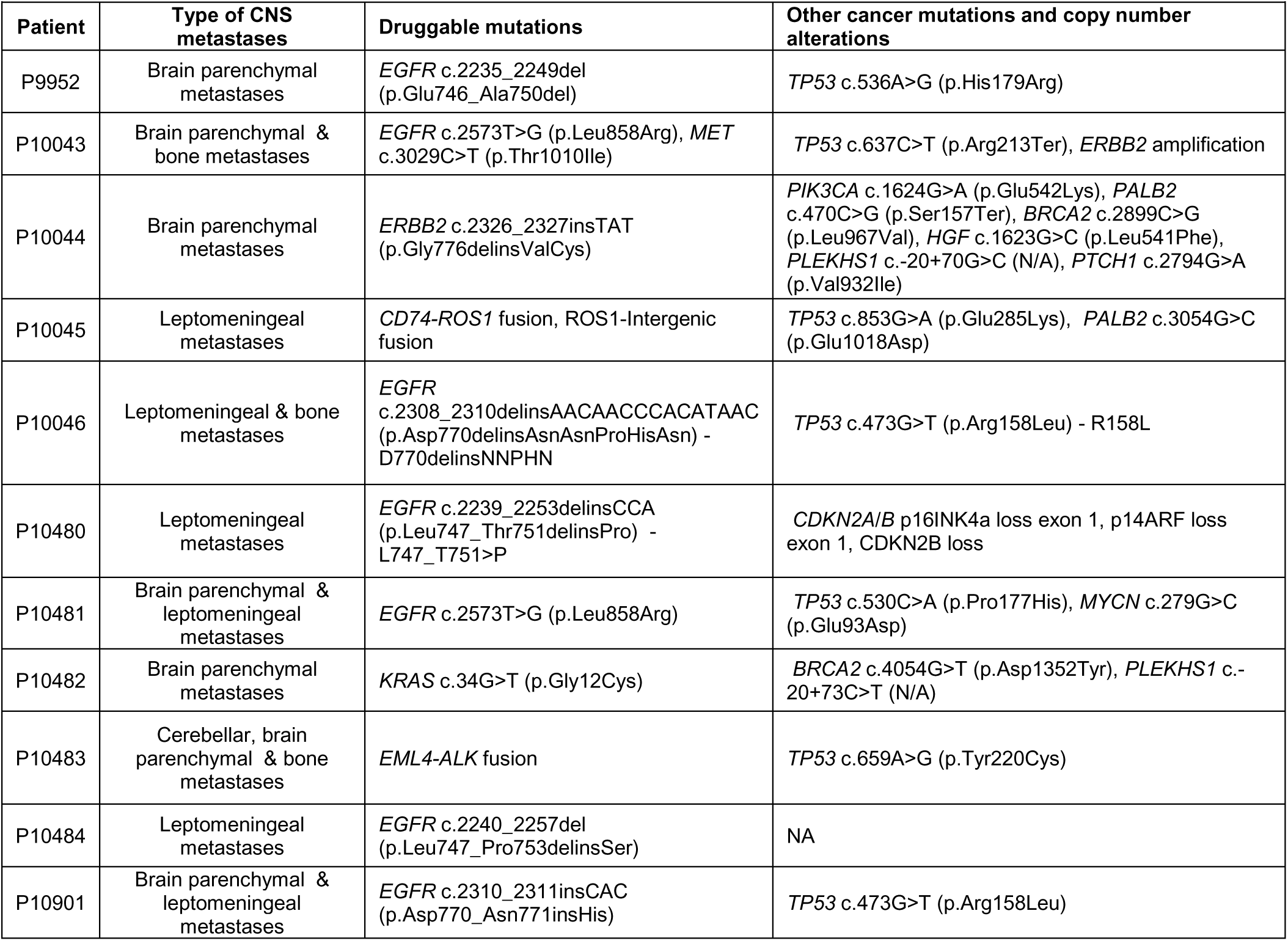
Clinical description of individuals with metastatic lung adenocarcinoma.

| **Patient** | **Type of CNS metastases** | **Druggable mutations** | **Other cancer mutations and copy number alterations** |
| --- | --- | --- | --- |
| P9952 | Brain parenchymal metastases | *EGFR* c.2235_2249del (p.Glu746_Ala750del) | *TP53* c.536A>G (p.His179Arg) |
| P10043 | Brain parenchymal & bone metastases | *EGFR* c.2573T>G (p.Leu858Arg), *MET* c.3029C>T (p.Thr1010Ile) | *TP53* c.637C>T (p.Arg213Ter), *ERBB2* amplification |
| P10044 | Brain parenchymal metastases | *ERBB2* c.2326_2327insTAT (p.Gly776delinsValCys) | *PIK3CA* c.1624G>A (p.Glu542Lys), *PALB2* c.470C>G (p.Ser157Ter), *BRCA2* c.2899C>G (p.Leu967Val), *HGF* c.1623G>C (p.Leu541Phe), *PLEKHS1* c.-20+70G>C (N/A), *PTCH1* c.2794G>A (p.Val932Ile) |
| P10045 | Leptomeningeal metastases | *CD74-ROS1* fusion, ROS1-Intergenic fusion | *TP53* c.853G>A (p.Glu285Lys), *PALB2* c.3054G>C (p.Glu1018Asp) |
| P10046 | Leptomeningeal & bone metastases | *EGFR* c.2308_2310delinsAACAACCCACATAAC (p.Asp770delinsAsnAsnProHisAsn) - D770delinsNNPHN | *TP53* c.473G>T (p.Arg158Leu) - R158L |
| P10480 | Leptomeningeal metastases | *EGFR* c.2239_2253delinsCCA (p.Leu747_Thr751delinsPro) - L747_T751>P | *CDKN2A/B* p16INK4a loss exon 1, p14ARF loss exon 1, *CDKN2B* loss |
| P10481 | Brain parenchymal & leptomeningeal metastases | *EGFR* c.2573T>G (p.Leu858Arg) | *TP53* c.530C>A (p.Pro177His), *MYCN* c.279G>C (p.Glu93Asp) |
| P10482 | Brain parenchymal metastases | *KRAS* c.34G>T (p.Gly12Cys) | *BRCA2* c.4054G>T (p.Asp1352Tyr), *PLEKHS1* c.-20+73C>T (N/A) |
| P10483 | Cerebellar, brain parenchymal & bone metastases | *EML4-ALK* fusion | *TP53* c.659A>G (p.Tyr220Cys) |
| P10484 | Leptomeningeal metastases | *EGFR* c.2240_2257del (p.Leu747_Pro753delinsSer) | NA |
| P10901 | Brain parenchymal & leptomeningeal metastases | *EGFR* c.2310_2311insCAC (p.Asp770_Asn771insHis) | *TP53* c.473G>T (p.Arg158Leu) |

### Features of brain metastasis CSF cfDNA

For each CSF sample, all isolated cfDNA was processed into a sequencing library – the entirety which was loaded into the nanopore sequencer (**Methods**). Sequencing metrics of all the samples are summarized in **SI Table 1**. As we had shown in a prior publication, the total number of mapping sequence reads correlated with the amount of DNA and its concentration [13]. We applied a normalization step to the sequencing data which accounted for the starting CSF volume from which cfDNA was isolated. This step provided a sequencing coverage metric that incorporated the volume of CSF used for each sample. Across the CSF cfDNA samples, the normalized sequencing coverage ranged from 0.01 to 7.0X, while the controls ranged from to 0.01 to 0.1X. We observed a general trend where the cancer patients had higher levels of cfDNA (**Figure 2A**). Notably, the cancer patients levels were significantly higher compared to non-cancer controls (*p*=0.0522) (**Figure 2B**). These results may be an indication that brain metastases release greater amount of DNA into the CSF compared to the normal homeostatic state of individuals without brain tumors.

**Figure 2.**
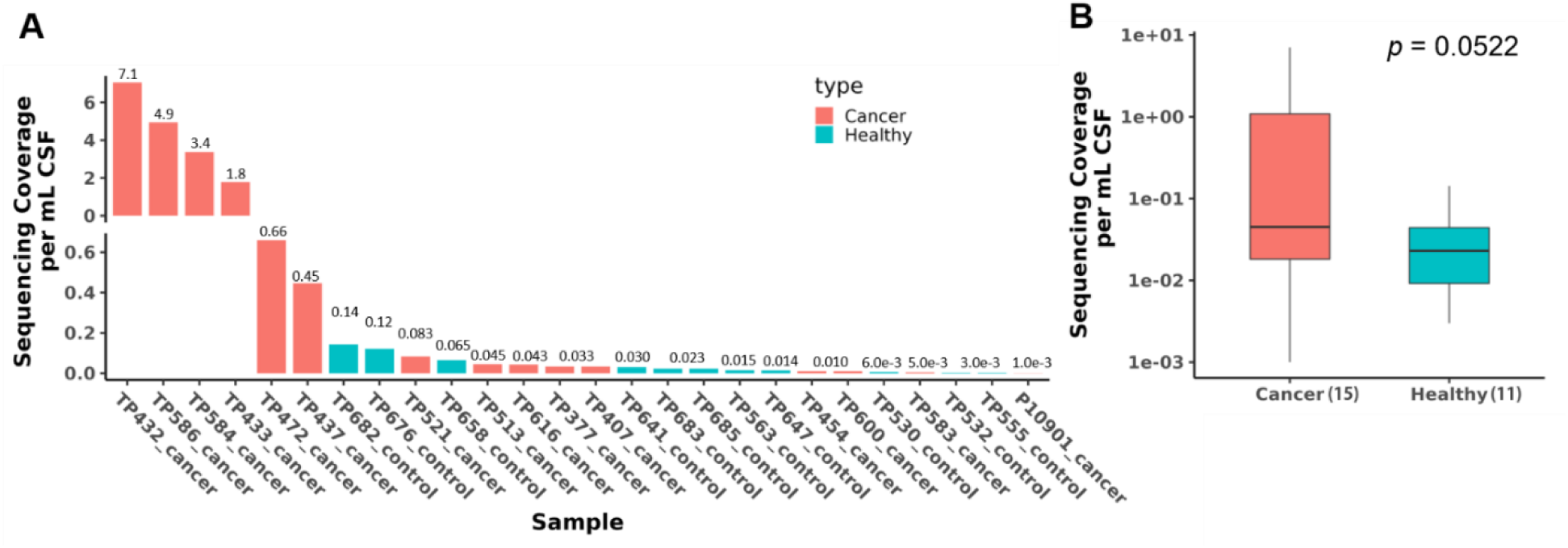
Analysis of sequencing coverage in 1 mL of CSF from NSCLC-BMET patients and healthy controls. Sequencing coverage were normalized to 1 mL of CSF for NSCLC-BMET patients (N=15) and non-cancer controls (N=11). B) Cancer sample showed higher sequencing coverage than controls (*p* = 0.0522). The normalized sequencing coverage ranged between 0.01-7X in cancer patients and 0.01-0.1X in controls with significant differences.

### Distinct fragmentation profiles of the CSF cfDNA of brain metastasis

We characterized the DNA fragment features from CSF cfDNA. Nanopore sequencing provides read lengths up to one Mb. This contrasts with short read sequencers which typically generate ones that are several hundred bases long. As a result, nanopore long reads can identify DNA from multiple nucleosome aggregates such as trinucleosomes which can exceed 400 bases in length. Overall, the length of the sequence read is directly reflective of whether the DNA length is organized as a mononucleosome, dinucleosome or trinucleosome.

We made comparisons were among four sources of the extracted cfDNA: (1) CSF from patients with metastatic NSCLC to the brain; (2) CSF from non-cancer controls; (3) plasma from patients with metastatic NSCLC to another organ site; (4) plasma from non-cancer controls. We show examples of the cfDNA fragment size distributions among non-cancer and cancer patients as shown in **SI Figure 1-3**.

**Figure 3.**
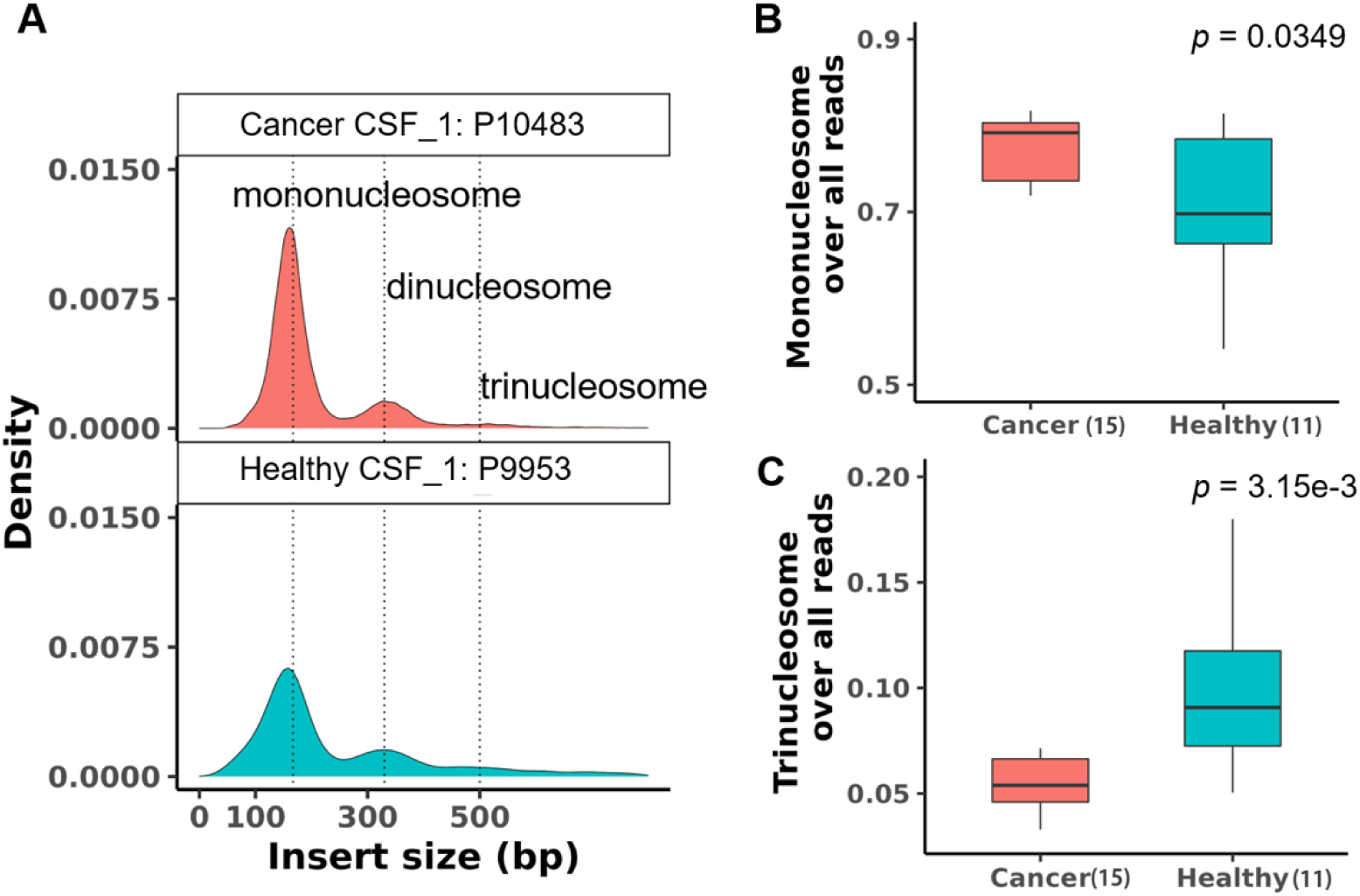
Analysis of CSF cfDNA mono-/trinucleosome ratios from NSCLC-BMET patients and healthy controls. Peaks of mono-nucleosome (167 bp), dinucleosome (330 bp) and trinucleosome (500 bp) were found in all samples. A) Examples of cfDNA fragment size distributions were shown for one cancer patient P10483 and one healthy control P9953. B) The overall fraction of each nucleosome type was calculated. B) The mononucleosome fraction was higher among cancer patients compared to healthy controls by a factor of 1.10 (*p* = 0.0349). C) The trinucleosome fraction was higher among the healthy controls compared to cancer patients by a factor of 1.81 (*p* = 3.15e-3).

We determined whether there were significant differences between NSCLC brain metastases compared to non-cancer controls. These fragments produced distinct peaks that correlated with the read lengths of 167bp, 330bp and 500 bp that were representative of mono-, di- and trinucleosomes, respectively (**Figure 3A**). We determined the overall fraction of each nucleosome type, specifically the number of each nucleosome size over the total. The mononucleosome fraction was higher among cancer patients compared to healthy controls by a factor of 1.10 (*p* = 0.0349) (**Figure 3B**). The trinucleosome fraction was higher among the healthy controls compared to cancer patients by a factor of 1.81 (*p* = 3.15e-3) (**Figure 3C**).

### Specific nucleosome ratios occur in the CSF cfDNA of brain metastasis

As we previously reported, the ratios of different size nucleosome fragments from cfDNA are highly informative for determining the presence of cancer [13]. For example, the plasma-derived cfDNA from colorectal cancer patients has a significantly higher mono-/dinucleosome ratio compared to cfDNA from healthy controls [13].

From the CSF cfDNA, we identified the mono-/dinucleosomes, mono-/trinucleosomes and di-/trinucleosome ratios for each sample and observed some general trends. NSCLC-BMET cancer patients had a higher mono-/trinucleosome ratio compared to controls (*p* = 7.31e-4) (**Figure 4A**). Similarly, cancer patients had a higher di-/trinucleosome ratio compared to controls (*p* = 6.95e-6) (**Figure 4B**). However, when comparing between cancer patients and controls, the mono-/dinucleosome ratios were not significantly different (*p* = 0.260) (**Figure 4C**). This result suggests that specific nucleosome fractions in the CSF are indicators of brain metastasis.

**Figure 4.**
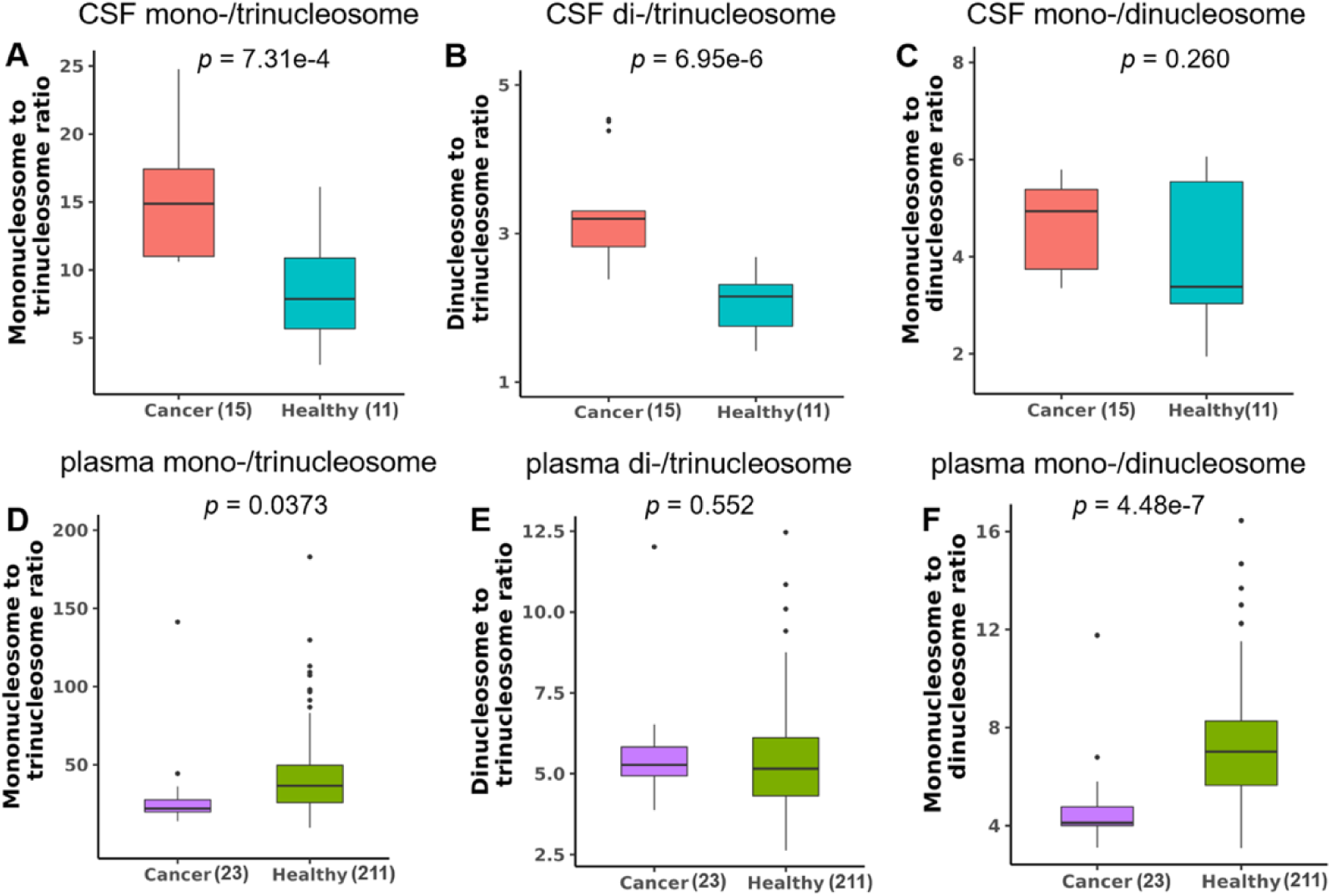
Analysis of cfDNA nucleosome ratios in CSF and plasma from NSCLC-BMET patients and healthy controls. The mono-/trinucleosome ratios (A) and di-/trinucleosome ratios (B) were significantly higher in cancer CSF (N=15) compared to control CSF (N=11). (C) The mono-/di-nucleosome ratios showed no significant difference between cancer CSF and control CSF. While in plasma, the patterns were completely different. The mono-/dinucleosome ratios (F) and mono-/trinucleosome ratios (D) showed significant differences between cancer plasma (N=23) and control plasma (N=211). The di-/trinucleosome ratios showed no significant difference between cancer plasma and control plasma.

### Comparing nucleosome ratios in plasma versus CSF cfDNA

Next, we sought to determine whether there are any differences in cfDNA fragmentation between different biological fluid sources. To do so, we compared the cfDNA fragmentation patterns derived from plasma versus those from CSF. For determining plasma cfDNA properties, we used an independent cohort of lung cancer patient and healthy control samples.

Sequencing metrics of all the plasma samples are summarized in **SI Table 2**. Comparisons and their p values are summarized in **SI Table 3** and **SI Figures 4**. The plasma samples came from patients with metastatic NSCLC in different organs including the brain. One general feature was that the plasma-derived cfDNA had significantly lower trinucleosome peak values compared to CSF derived cfDNA (*p* = 3.08e-7) (**SI Figure 5A**). This was the case for both the metastatic NSCLC patients (*p* = 3.74e-5) and healthy controls (*p* = 8.37e-5) (**SI Figure 5B**).

**Figure 5.**
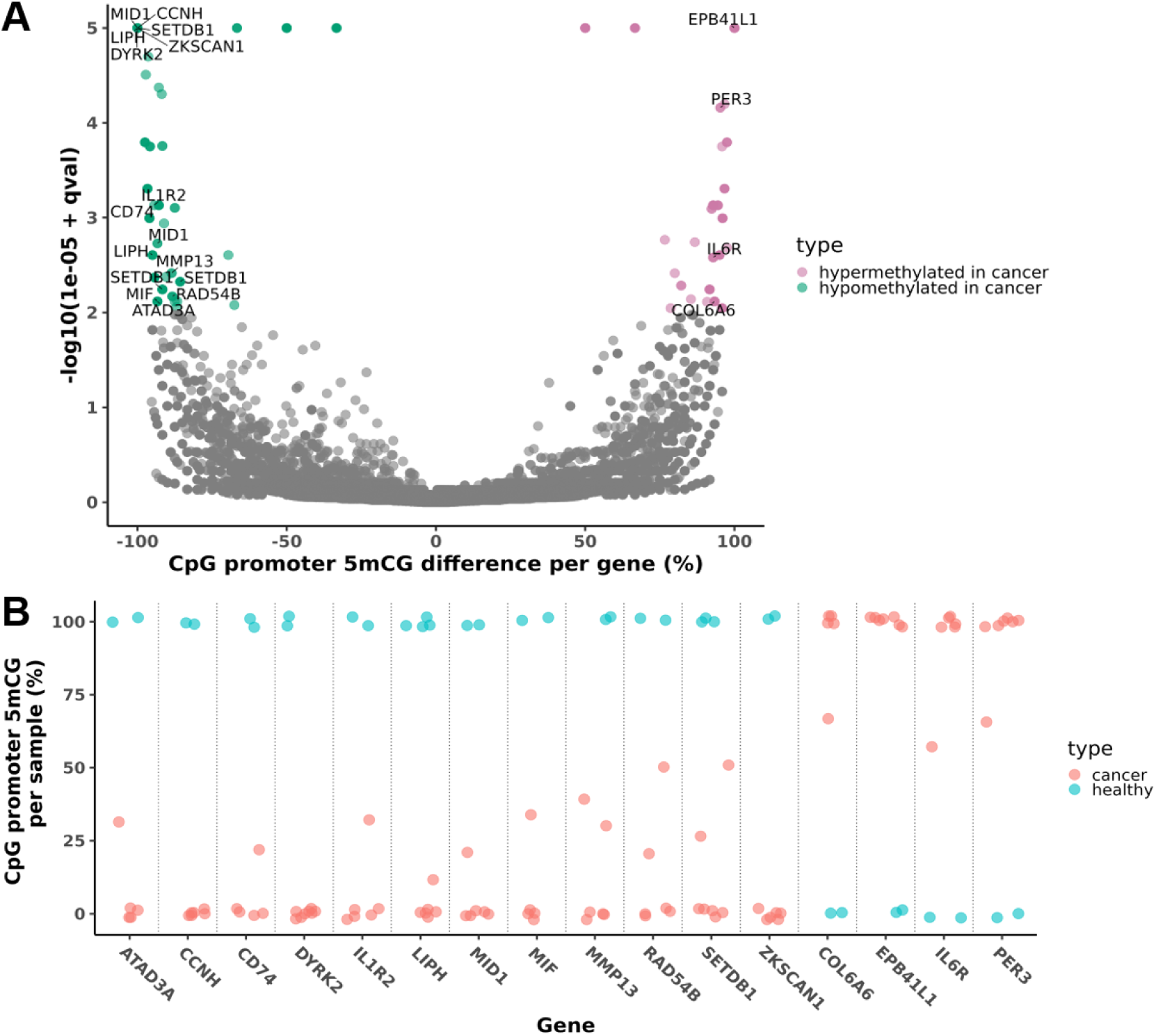
Analysis of methylated cfDNA for promoter 5mCG in cancer patients. A) Genes with differential methylated between cancer vs. control on promoter CpG sites were plotted. Each dot represents gene(s) with same FDR-corrected q value. Colored dot were genes at q value < 0.01 with hyper-/hypomethylation in cancer vs. control. CpG-site 5mCG methylation levels of 16 reported biomarkers from 176 differentially methylated promoter genes were plotted. Each dot represents a sample. Distinct methylation patterns were shown between cancer patients and healthy controls.

When examining the cfDNA nucleosome ratios in plasma compared to CSF, we observed several striking differences. For the plasma cfDNA from NSCLC patient compared to controls, there was borderline statistical significance for the mono-/trinucleosome ratios (*p* = 0.0373) (**Figure 4D**) and no significant difference for the di-/trinucleosome ratios (*p* = 0.552) (**Figure 4E**). However, there was a significantly lower mono-/dinucleosome ratio compared to the non-cancer controls (*p* = 4.48e-7) (**Figure 4F**). This difference in the plasma cfDNA nucleosome ratios contrasts with what was observed in CSF cfDNA from BMET patients. Specifically, for CSF cfDNA from NSCLC patients with BMETs compared to controls, there was a significantly increased mono-/trinucleosome and di-/trinucleosome ratios (**Figure 4A-B**). Overall, these results indicate that the plasma cfDNA fragmentation has lower trinucleosome content compared to the CSF cfDNA.

### Specific 5mC methylation patterns present in the CSF cfDNA of brain metastasis

Nanopore sequencing of CSF cfDNA directly identifies 5mC methylation. It does so without any chemical or biochemical conversion processes and PCR amplification. We conducted an analysis on the CSF cfDNA’s 5mC methylation patterns, comparing the NSCLC-BMET patients versus non-cancer controls. We determined the differential 5mC alterations among CpG promoter sites - the promoter regions were defined as being 2kb upstream and 500bp downstream of the transcription start site. After this step we identified the genes with differentially methylated CpG sites per FDR-corrected q values. These results are shown on a volcano plot with q values where each dot represents a gene (**Figure 5A**). Next, we applied a threshold to identify those differentially methylation sites present in six or more samples and having a q value < 0.01. Based on this ranked criteria, there was differential methylation of 176 promoter sites for protein-encoding genes (**SI Table 4**).

Among the 176 genes, we determined which ones had functional significance in NSCLC. For this analysis, we conducted a literature review for NSCLC studies reporting increased versus decreased methylation for specific genes, increased versus decreased tumor specific gene expression biomarkers and involvement in specific cancer pathways. We identified 16 genes with evidence for a functional role in NSCLC. These 16 gene markers are labeled on the volcano plot in **Figure 5A** and notably, all were among the most statistically significant in the volcano plot. Among the 16, there were 12 genes with decreased methylation. These genes were reported to have upregulated gene expression, as observed in NSCLC studies and included *ATAD3*, *CCNH*, *CD74*, *DYRK2*, *IL1R2*, *LIPH*, *MID1*, *MIF*, *MMP13*, *RAD54B*, *SETDB1* and *ZKSCAN1*. Citing an example, the increased expression of *ZKSCAN1* promotes NSCLC progression by inactivating MAPK signaling pathway [22]. Increased *SETDB1* expression stimulates the Wnt/β-Catenin pathway and decreased *TP53* expression in NSCLC [23].

There were four genes with hypermethylation. These genes had been previously described as having downregulated gene expression in NSCLC and included *COL6A6*, *EBP41L1*, *IL6R* and *PER3*. Yang et al. reported that *EPB41L1* hypermethylation in NSCLC [24]. A *COL6A6* knockout in a NSCLC cell line was reported to increase the rate metastasis by activating the JAK signaling pathway [25].

Next, we determined which CSF samples had measurable promoter methylation among these 16 genes. This analysis involved quantifying the methylated promoter CpG sites per a given gene versus non-methylated sites for both the cancer and control CSF samples (**Figure 5B**). There is a clear trend that these genes were highly informative for distinguishing cancer versus normal CSF samples. For decreased methylation, all cancer samples showed extensive degree of promoter methylation. When considering the promoters of these 16 genes, 80% (12/15) of cancer samples and 91% (10/11) of the non- cancer controls had differential methylation. This result indicated that these genes methylation status is present among multiple samples.

Based on the 176 genes with differential methylation, we determined which biological pathways and protein-protein interactions (**PPI**) were associated with these genes. We used the Enrichr algorithm to identify these broader features [26]. The pathway analysis showed functional pathways associated with *EGFR* mutation positive NSCLC. These pathways included cell signaling via IGF1, PI3K/TRK, PTP1B, c-Kit receptor and cell cycle signaling (p<0.012) (**Figure 6A**). The *EGFR* mutation leads to the activation of PI3K/Akt/PTEN/mTOR and MAPK pathways. PTP1B activates the ERK1/2 and modulates the MAPK pathway [27]. Protein-protein interaction (**PPI**) partners included PRKACA, CASP3, SRC, JAK2 (p<0.05); these genes are involved in the progression of NSCLC. Citing an example of biological pathway interactions, SRC activates EGFR and the downstream pathways like MAPK [28]. The JAK2/STAT3 pathway is activated in EGFR-mutant NSCLC [29] (**Figure 6B**).

**Figure 6.**
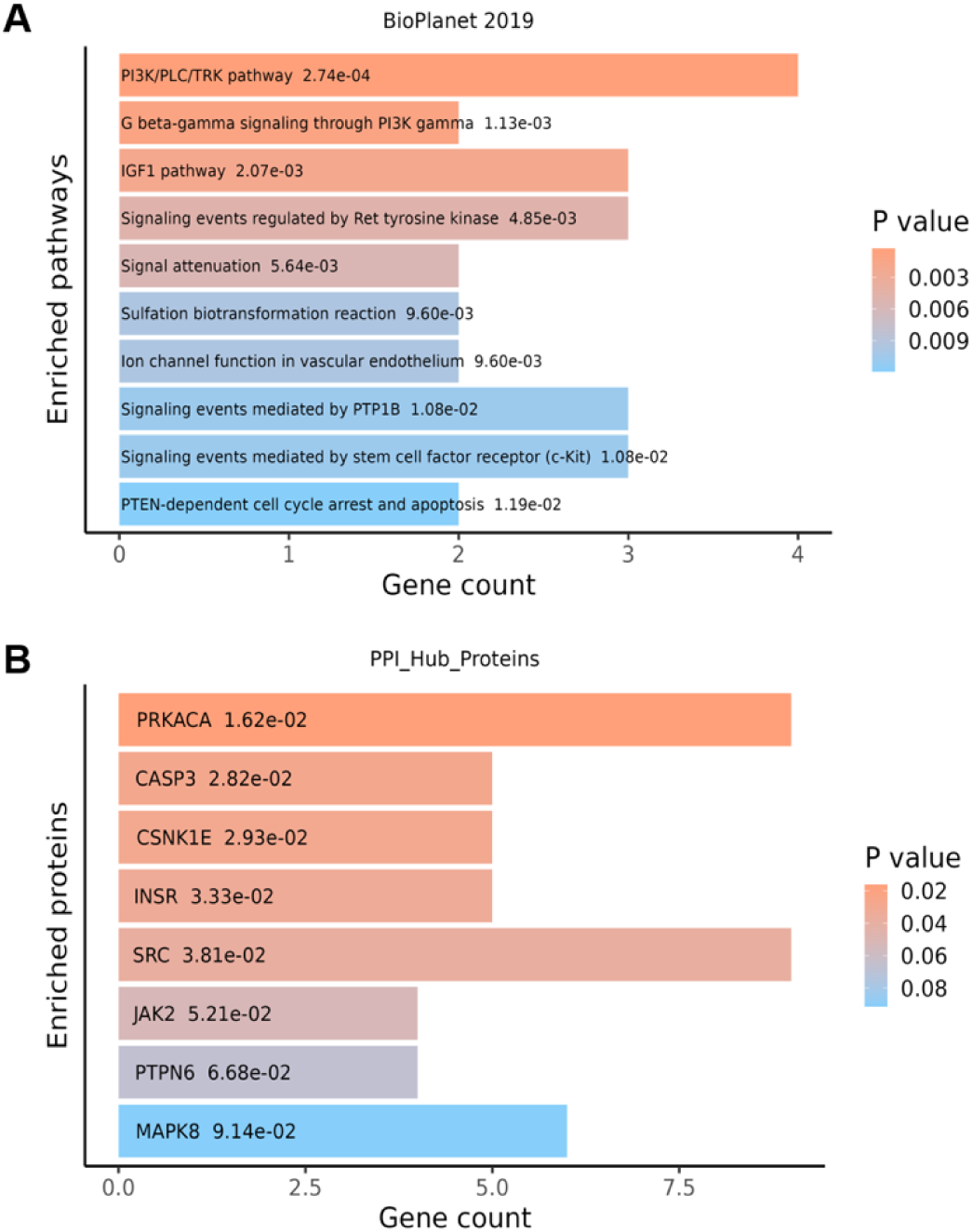
Analysis of enriched pathways and protein-protein interactions for promoter 5mCG in cancer patients. A) Pathway enrichment analysis was performed by Enrichr algorithm. The p-values were shown for each associated pathway. B) Protein-protein interaction analysis was performed with the PPI Hub Proteins database. The p-values are shown for each associated protein.

A subset of patients had NSCLC *EGFR* mutations (exon 19 deletion, L858R mutation and exon 20 insertion). We further evaluated the CSF cfDNA differential methylation among these patients. Most CSF samples had differential methylation in seven genes, specifically *DYRK2*, *LIPH, SETDB1, ZKSCAN1, EBP41L1, IL6R* and *PER3*.

### Specific 5hmC hydroxymethylation patterns in the CSF cfDNA of brain metastasis

Nanopore sequencing of CSF cfDNA directly identifies hydroxymethylation at specific sites without the need for additional molecular processing. The 5hmC pattern has a different distribution than 5mC. Hydroxymethylation occurs more frequently in both a gene’s exon and intron sequence [30]. Moreover, increased levels of intragenic hydroxymethylation (5hmC) are positively correlated with increased gene expression, more so than intragenic methylation (5mC) changes [30, 31]. Therefore, we evaluated the differential hydroxymethylation changes in the intragenic gene body and promoter sequences.

Decreased hydroxymethyation was observed across all methylated CpG sites per a given sample (**SI Table 1**). Notably, cancer CSF cfDNA had a significantly lower average genome-wide 5hmC% (*p* = 8.31e-4) compared to the controls (**Figure 7**). In contrast, there was no increased hydroxymethylation among the cancer samples.

**Figure 7.**
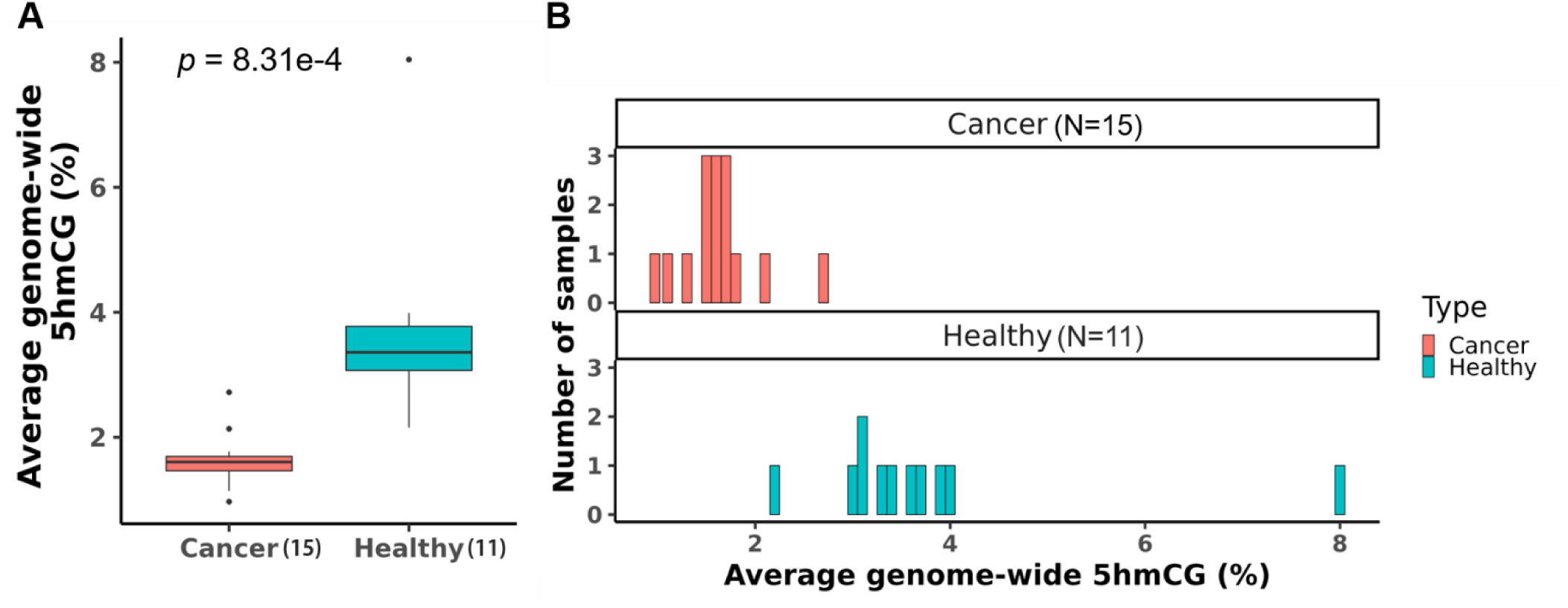
Analysis of average genome-wide 5hmCG hydroxymethylation of NSCLC-BMET patients and healthy controls. The genome-wide average 5hmCG% were significantly lower in cancer patients than healthy controls. Data plotted in A) box plot and B) bar plot.

### Decreased intragenic hydroxymethylation in the CSF cfDNA of brain metastasis

Comparing the NSCLC-BMET versus the non-cancer CSF cfDNA, we identified the genes with differential 5hmC intragenic sites. These results are shown on a volcano plot with each dot representing a gene (**Figure 8A**). We calculated the intragenic 5hmC differences as percentages. We applied a sample threshold such that a differentially hydroxymethylated site was present in eight or more samples (q value < 0.01). We focused on the top ranked genes with the most decrease in hydroxymethylation for the cancer samples compared to controls.

**Figure 8.**
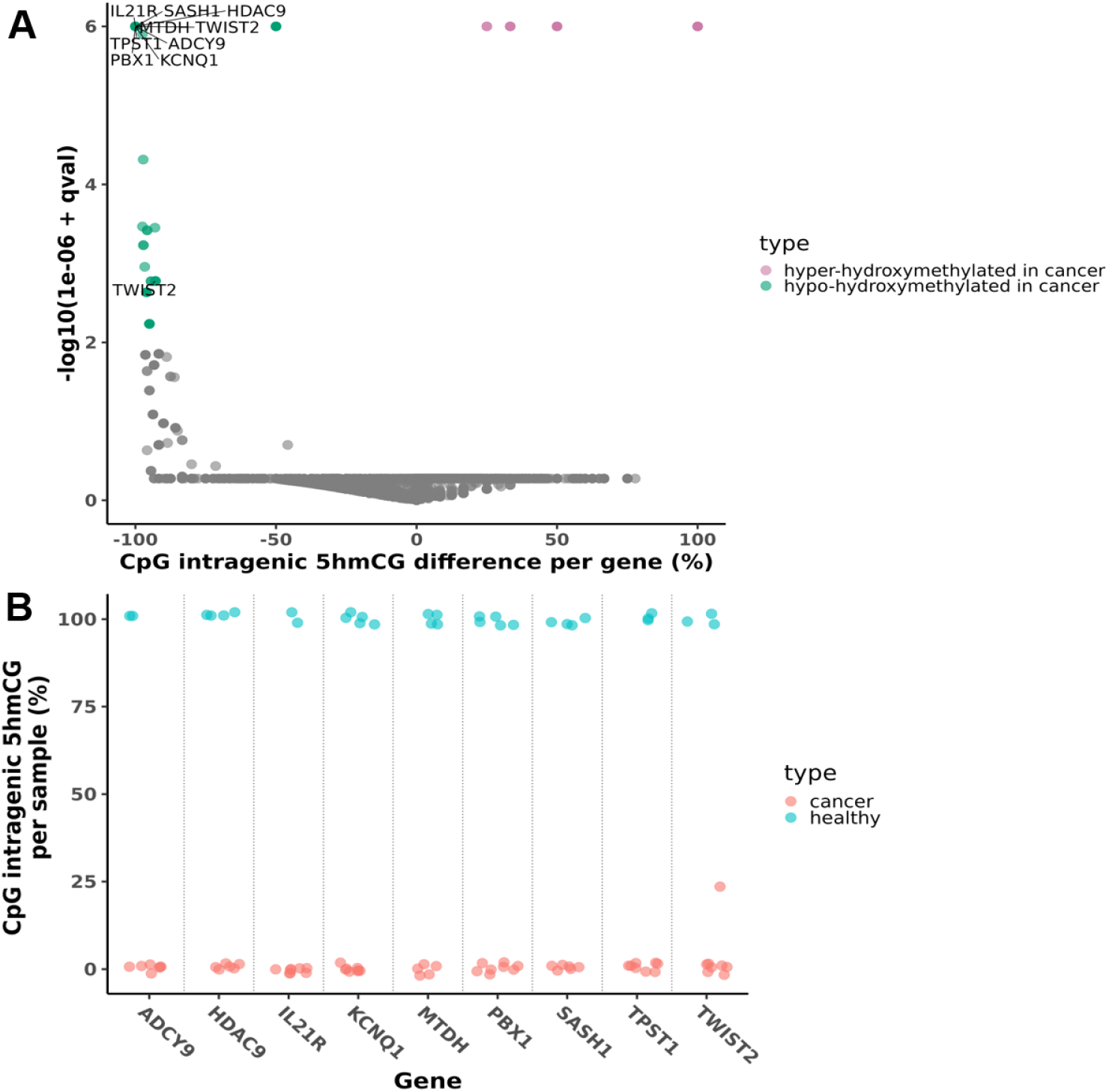
Analysis of methylated cfDNA for intragenic 5hmCG in cancer patients. A) Genes with differential hydroxymethylated between cancer vs. control on intragenic CpG sites were plotted. Each dot represents gene(s) with same FDR-corrected q value. Colored dot were genes at q value < 0.01 with hyper-/hypo-hydroxymethylation in cancer vs. control. B) CpG-site 5hmCG methylation levels of 9 reported biomarkers from 109 differentially hydroxymethylated intragenic genes were plotted. Each dot represents a sample. Distinct methylation patterns were shown between cancer patients and healthy controls.

Based on these criteria, there were 109 genes with significantly decreased hydroxymethylation (**SI Table 4**).

Among the top 109 genes, we determined which ones had functional relationship to NSCLC. We identified published reports describing NSCLC genes with reduced hydroxymethylation, altered gene expression and involvement in cancer biological pathways (**Methods**). As noted in the NSCLC literature, there were nine genes with intragenic decreased hydroxymethylation, correlated downregulated gene expression or tumor suppressor function. The genes associated with NSCLC are color labeled on the volcano plot in **Figure 8A**. There was a total of 14 genes with decreased hydroxymethylation changes. Next, we determined the number of hydroxymethylated intragenic CpG sites over the total for each of these genes in both cancer and control samples (**Figure 8B**). Each dot represents a CSF sample. Among the 14 genes, hydroxymethylation changes were observed in 88% (23/26) of all the samples, with 80% (12/15) in cancer and 100% (11/11) among the controls.

Nine genes had decreased hydroxymethylation. They were *ADCY9*, *HDAC9*, *IL21R*, *KCNQ1*, *MTDH*, *PBX1, SASH1*, *TPST1* and *TWIST2* (**Figure 8B**). Citing some examples of their functional relationship in NSCLC, the IL21/IL21R axis was reported to reduce cancer cell growth by inhibiting Wnt/β-Catenin signaling and PD-L1 expression [32]. The *SASH1* gene is a tumor suppressor and may be a prognostic indicator in NSCLC [33].

There were five genes that showed increased hydroxymethylation albeit in fewer samples (i.e., five). Among them, two genes, *CIZ1* and *PRRG4*, showed increase expression in NSCLC. The *CIZ1* gene (b variant) was reported as a circulating biomarker for early NSCLC [34]. The expression was high both in plasma and tissue from NSCLC patients. Another report showed that the long non-coding RNA PRRG4-4 was upregulated in NSCLC tissues [35].

Using the Enrichr algorithm, we conducted pathway analysis and protein-protein interaction analysis on the 109 genes. The pathway analysis showed increase among specific biological pathways related to NSCLC. They included endothelins, Ras activation, and the estrogen receptor (**ER**) (p<0.001) (**Figure 9A**). Inhibition of EGFR with a mutation induces the secretion of endothelins [36]. ERs are reported to activate EGFR in a reciprocally. ERβ activates downstream substrates of EGFR like PI3K and MAPK [37].

**Figure 9.**
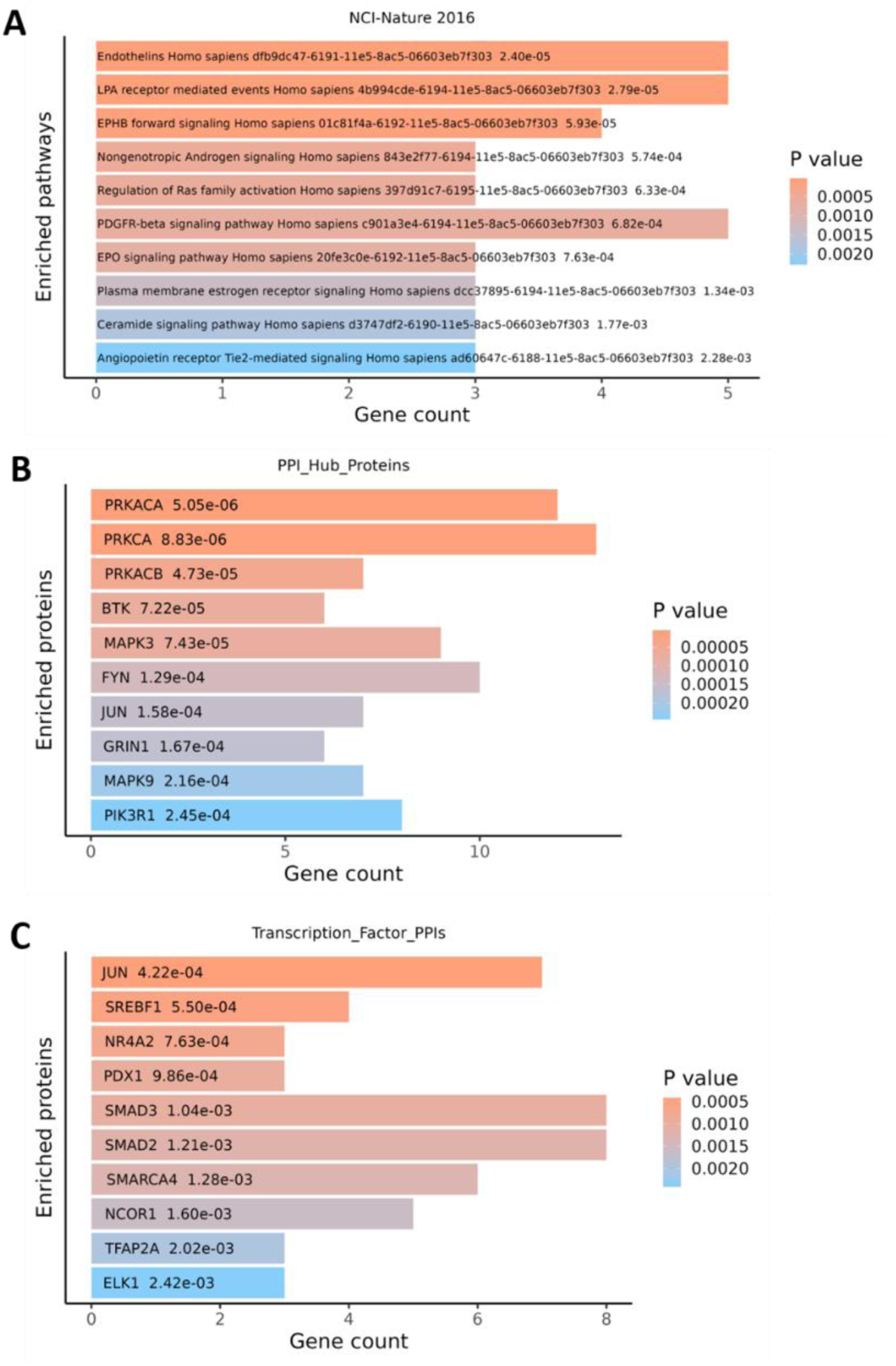
Analysis of enriched pathways and protein-protein interactions for intragenic 5hmCG in cancer patients. A) Pathway enrichment analysis was performed by Enrichr. The p-values were shown for each associated pathway. B) Protein-protein interaction analysis was performed by Enrichr using PPI Hub Proteins database. C) Protein-protein interaction analysis was performed by Enrichr using Transcription Factor PPIs database. p-values were shown for each associated protein.

For these 106 genes, protein-protein interaction (**PPI**) partners included PRKACA, PRKACB, PRKCA, BTK, MAPK3, PIK3R1 (*p* <2.5e-4) (**Figure 9B**), SMAD2, SMAD3, SMARCA4, NCOR1, TFAP2A, ELK1 (*p* <2.4e-3) (**Figure 9C**). All proteins were involved in the progression of NSCLC. Citing some specific examples in NSCLC cell lines, when a BTK inhibitor suppresses the STAT3/JAK2/Akt axis there is a potentiation of the effects of EGFR-TKIs treatment [38].

PI3KR1 is involved in the PI3K/Akt/mTOR pathway in EGFR-mutant NSCLC. SMAD2 and SMAD3 are key elements in the downstream of TGF-β signaling pathway in NSCLC [39]. TFAP2A activates MAPK pathway in NSCLC [40]. This signaling pathway contributes to EGFR-TKI resistance and the silencing of ELK1 improves TKI killing in cancer cell lines [41].

### Decreased promoter hydroxymethylation in the CSF cfDNA of brain metastasis

We also identified 89 genes with decreased hydroxymethylation in the promoter region and conducted a pathway analysis on these genes. Among the BMET samples, the promoter 5hmC changes were enriched with immune response signaling pathways which included the IFN-γ, IL- 3 and IL-5. Oher cellular signaling pathways included PDGFβ, MAPK, and ERBB1 (p<0.02) (**SI Figure 6A**). The PPI analysis showed associated proteins like CDK2, CREB1, HDAC4, RAC1, APC and PIK3R1 (p<0.05) (**SI Figure 6B**). Others with lower statistical significance include FOXP3, KDM6A, ELK4, ZEB1, FLI1 and STAT3 (p<0.1) (**SI Figure 6C**). These proteins are also involved in the progression of NSCLC.

### Comparing the methylation of brain metastasis CSF cfDNA to NSCLC primary tumor DNA

We compared the differentially methylated (5mC) genes from the CSF cfDNA versus those genes reported in the TCGA-LUAD genomic data set which comes from NSCLCs. For this analysis, we used the CanMethdb [20]. This resource provides methylation status of CpG sites from TCGA data sets and others. It includes the upstream cis-regulatory elements as well as functional annotations for the target genes. We cross referenced differentially methylated sites in the intragenic and promoter region (**Methods**). The two promoter annotations were based on the following criteria: (1) an interval that covered 2 kb upstream and 500 bp downstream of transcription start site from GENCODE v38, and (2) a promoter annotation using Ensembl (release 112). The results are summarized in **SI Table 5-6**. We identified the differentially methylated gene promoters that overlapped between the two data sets. There were 48 (17%) with decreased methylation and 60 (16%) with increased methylation. For decreased methylation, six genes have been reported as NSCLC biomarker: *ATAD3A* [42], *MMP13* [43], *DRYK2* [44], *NQO1* [45], *MIF* [46] and *RAD54B* [47]. For those genes with increased methylation, one was a reported NSCLC biomarker as *COL6A6* [25].

To characterize chromatin accessibility among the TCGA NSCLC samples, we analyzed the ATAC-seq datasets of lung tumors [21]. This assay identifies accessible chromatic regions. Then we identified the methylated promoters, as found in the CSF cfDNA analysis, which overlapped with open chromatin sites per the ATAC assays as listed in the TCGA NSCLC study. The results are summarized in **SI Table 7**. Comparing the overlapping methylated gene promoters versus open chromatin sites, there were 65 (17%) with decreased methylation and 36 (7%) with increased methylation. Among the genes with decreased methylation, six were previously reported as NSCLC biomarkers*: SETDB1* [23], *MMP13* [43], *DYRK2* [44], *MIF* [46], *LIPH* [48] and *MID1* [49].

### Intra-individual variation is stable in CSF cfDNA profiles

This study had samples from four patients with two sequential draws at different time intervals. Three patients had short time intervals over 7-21 days with no evidence of clinical progression or changes (P9952, 10043, 10483, Group 1). One patient had two samples that were obtained over 288 days (P10481, Group 2). For the paired samples, we compared three features: the nucleosome ratios, average genome-wide 5mC%, and average genome-wide 5hmC (**SI Table 8**). We determined the percentage change between the paired samples. For the Group 1, the three features had a low percentage change (<20%). These patients had stable CNS disease with no evidence of progression and clinical course. This result suggests that the CSF cfDNA profile was stable. The Group 2 patient (P10481) showed decreased mono-/di- and mono-/tri-nucleosome ratios with percentage changes at 35.3% and 37.4%, respectively. This variation was related to different treatments including radiation.

## DISCUSSION

We conducted a multi-omic analysis of CSF-derived cfDNA from patients with NSCLC BMETs. This study leveraged nanopore sequencing to evaluate fragmentation, methylation (5mC) and hydroxymethylation (5hmC) from the same cfDNA molecules. Among patients with BMET, we discovered that the CSF cfDNA had distinct genomic features not observed in CSF cfDNA from healthy controls. These BMET features included DNA fragmentation shifts in the mono-/trinucleosome ratios, specific methylation patterns and distinct hydroxymethylation patterns. Overall, this is the first work to identify distinct fragmentation profiles of mono-, di- and trinucleosomes in CSF-derived cfDNA from cancer patients. These distinct features such as CSF cfDNA nucleosome ratios may be useful biomarkers for identifying the presence of NSCLC-BMET.

Many studies in cfDNA fragmentation have used short read sequencing to focus on fragment size less than 250 bp which are mono-nucleosomes. However, our study showed the advantages of nanopore sequencing which provides longer reads. Our results are in line with another report in which nanopore sequencing provided longer cfDNA fragments greater than 300 bp compared to short read sequencing (i.e., Illumina) in plasma and urine [50]. Additionally, a prior article from our group showed the enriched mono-nucleosome patterns in plasma-derived cfDNA from colorectal cancer patients based on mono-/dinucleosome ratios [13]. We applied the same nanopore sequencing approach to CSF-derived cfDNA. We observed trinucleosome peaks directly from the fragment size distributions, which were absent for plasma-derived cfDNA, either from cancer patients or healthy controls, analyzed with the same approach. We also observed distinct mono-/trinucleosome ratios in CSF-derived cfDNA from NSCLC-BMET patients in our cohort. This fragmentation pattern may indicate higher cfDNA fragmentation in CSF than plasma. This study provided a comprehensive characterization of cfDNA fragmentation patterns in CSF-derived cfDNA from NSCLC-BMET patients.

There have been a variety of cancer studies on plasma-derived cfDNA and their fragment patterns [51]. For example, one study noted that there were smaller mono-nucleosome fragments with a size of ∼143 bp compared to healthy controls with a size of ∼167bp [42]. Another study used a genome-wide approach to distinguish fragment size differences between cancer and healthy cfDNA samples and determined the ratio of small cfDNA fragments (100- 150 bp) to larger cfDNA fragments (151-220 bp) [52]. However, there are very few studies on the cfDNA fragmentation were reported in CSF, especially the characterization of trinucleosomes. In a study of gliomas which a tumor originating in the brain, Mouliere et al. showed that there was enrichment of short DNA fragments at ∼145bp in CSF-derived cfDNA compared to matched plasma samples [53]. Another group reported similar observations from NSCLC patients with leptomeningeal metastases [54]. White et al. found compared the ratio of cfDNA fragments (140-200 bp) to fragments (300-360 bp) between CSF and plasma from patients with leptomeningeal disease [55]. The ratio was significantly higher in plasma-derived cfDNA than CSF-derived cfDNA.

Bisulfite sequencing has been used to identify 5mC methylation from CSF-derived cfDNA in NSCLC BMETs [48]. There is a study of cfDNA 5mC methylation in CSF of BMET from NSCLC [56]. This article reported a limited number of 5mC methylation sites in primary lung tumors that differentiated patients with BMETs.

Hydroxymethylation is recognized as a source of epigenetic regulation in cancer. The inverse relationship between 5hmC content and cell division rate is present in rapidly dividing cells.

Exceptions include embryonic stem cells, highly proliferating cancer cell lines and tumors wherein no or an extremely low levels of 5-hmC are detectable [57]. Regarding characterization of cfDNA 5hmC hydroxymethylation profiles in cancer, most studies used affinity-based methods like hMe-Seal [58] to selectively enrich 5hmC-containing DNA fragments for amplification and sequencing. This sequencing enrichment assays is limited to sequencing fragments from 5hmC and does not provide exact base pair resolution of the hydroxymethylation status. Compared to these methods, nanopore sequencing approach does not rely on 5hmC isolation or enrichment. Importantly, nanopore sequencing and its downstream base calling directly provide 5mC and 5hmC sites from native single molecules without other experimental assays. To this date, there are no reports on cfDNA 5hmC profiling using nanopore sequencing.

We reported for the first time of the different 5hmC levels, enriched pathways, and biomarkers in CSF-derived cfDNA from NSCLC-BMET patients and healthy controls. In this study, CSF cfDNA samples had a range of 1-8% CpG sites being hydroxymethylated while 46-73% CpG sites were methylated. Regardless of the small percentage of 5hmC in all CpG sites, the cancer samples showed significantly lower average genome-wide 5hmC% than healthy controls. Other studies have observed a general loss of 5hmC in plasma-derived cfDNA and tumor tissues from metastatic lung cancer [58, 59].

We also determined that the hydroxymethylation patterns from intragenic versus promoters identified different pathway activities. In the intragenic 5hmC, analysis of biomarkers and protein-protein interaction showed tumor suppressor genes and related proteins in cancer progression like PRKACA. We identified enriched pathways like regulation of Ras family – activation of this pathway enhances cancer cell proliferation and migration. However, in the promoter region, protein-protein interaction showed immune-related proteins like CDK2 [60] and FOXP3 [61]. We identified enriched pathways such as IFN-γ, IL-3 and IL-5 signaling pathways. These activated immune signals may indicate the breakdown of blood barriers and alternation of immune homeostasis in CSF [62]. Other studies have reported plasma-derived 5hmC enrichment in immune activation related genes in NSCLC patients that respond to immunotherapies [63].

Some samples were from patients with NSCLC *EGFR* mutations. Tyrosine kinase inhibitors (**TKIs**) such as osimertinib are effective treatment for NSCLC-BMET patients with *EGFR* mutations. However, extended treatment eventually leads to therapeutic resistance. Some of the marker genes from our results have a relationship with EGFR-TKI treatment. For example, in the case of promoter methylation, *NECTIN4* was hypomethylated in *EGFR* mutation positive NSCLC CSF cfDNA. This gene has been reported as an upregulated biomarker of NSCLC. Patients with low NECTIN4 level showed better survival than patients with high NECTIN4 level after osimertinib treatment [64]. The gene *CD24* was hypomethylated in *EGFR* mutation positive NSCLC. There is a report in the literature of increased CD24 expression after EGFR-TKI treatment which is an indicator of resistance [65]. Among our samples, the intragenic hydroxymethylation of *DNMT3A* was reduced in *EGFR* mutation positive NSCLC and increased in *EGFR* mutation negative NSCLCs. One study reported that *DNMT3A* mutations were lower in icotinib/gefitinib-resistant patients than drug-sensitive patients [66]. There were also markers found to regulate EGFR pathways. Details of these genes and their roles were summarized in **SI Table 9-10**.

We identified overlapping genes between our methylation data and TCGA NSCLC 450K data. Among them, 13 hypermethylated and 20 hypomethylated genes were also found in TCGA-LUAD. This result suggests that the CSF cfDNA methylation profile reflects the presence of NSCLC BMET. Seven genes, including one hypermethylated gene *COL6A6* and 6 hypomethylated genes (i.e., *ATAD3A, MMP13, DYRK2, NQO1, MIF, RAD54B*) were reported in literature as biomarkers for NSCLC. For example, COL6A6 knockout was found to accelerate the metastasis of NSCLC by activating the JAK signaling pathway [25]. We also studied the overlap between hypomethylated promoter genes from our data and open chromatin sites from the TCGA NSCLC data. We found 26 overlapping hypomethylated gene promoters. Six of the related genes (i.e., *SETDB1*, *MMP13, DYRK2, MIF, LIPH, MID1*) were reported in literature as upregulated biomarkers for NSCLC. For example, SETDB1 positively stimulated Wnt/β-Catenin pathway and decreased P53 expression [23].

## CONCLUSIONS

This study is the first report characterizing the NSCLC-specific fragmentation and hydroxymethylation profiles in CSF-derived cfDNA. We applied a single-molecule sequencing method based on nanopore sequencing to study the fragmentation, methylation and hydroxymethylation of CSF-derived cfDNA in NSCLC-BMET patients. We found significantly enriched mono-nucleosome patterns in cancer patients compared to healthy controls – specifically, this was observed in the mono-/trinucleosome ratio values. The comparison of the CSF- versus plasma-derived cfDNA showed that this informative mono-/trinucleosome ratio is specific to CSF. Leveraging nanopore sequencing, we directly obtained 5mC and 5hmC molecular information from the same set of DNA molecules and our analysis showed distinct methylation or hydroxymethylation patterns between cancer and control. Cancer patients showed significantly lower genome-wide average 5hmC% compared to healthy controls. We also observed different 5hmC profiles between intragenic and promoter regions. For the intragenic 5hmC markers, related pathways were enriched in cancer progression. In contrast, the promoter 5hmC pointed to immune response pathways. We anticipate the profiles of fragmentation, methylation and hydroxymethylation may prove to be useful for the early detection of NSCLC-BMET with liquid biopsies from CSF. Extended sample cohort are required to further validate our findings.

## List of abbreviations

cfDNA: Cell-free DNA
ctDNA: Circulating tumor
DNA NSCLC: Non-small cell lung cancer
BMET: Brain metastasis
CSF: Cerebrospinal fluids
5mC: 5-Methylcytosine
5hmC: 5-Hydroxymethylcytosine

## DECLARATIONS

### Ethics approval and consent to participate

We obtained informed consent from all patients based on a protocol (12625, 62499) approved by Stanford University’s Institutional Review Board. This study was conducted in compliance with the Helsinki Declaration.

### Consent for publication

Not applicable

### Availability of data and materials

Sequence-aligned BAM files and associated methylation calls are deposited in NCBI’s dbGaP under accession phs003794. Scripts related to the study are available at the URL listed below: https://github.com/Tianqi-Kiki/nanopore_CSF_cfDNA.

### Competing interests

The authors declare that they have no competing interests.

### Funding

The authors acknowledge financial support from the Clayville Foundation (to H.P.J.), NIH NCI (U54CA261717 to H.P.J., G.B., T.T.H.T., T.C. and M.H.G; U01CA282212 to B.T.L., X.B. and H.P.J.), NIH NHGRI (R35HG011292 to B.T.L. and T.C.), and Stanford Medicine Translational Research and Applied Medicine (TRAM) pilot grant (to T.C.).

### Authors’ contributions

T.C., B.T.L., and H.P.J. conceived the study. M.H.G established and oversaw the translational patient recruitment component of the study. G.B. and T.T.H.T. processed the samples. T .C. performed cfDNA extraction and the sequencing experiments. T.C., X.B., B.T.L., and H.P.J. analyzed the data. T.C., and H.P.J. wrote the manuscript. All authors reviewed and approved the final manuscript.

## Acknowledgements

Figures 1 is illustrated using BioRender.

## Notes

https://github.com/Tianqi-Kiki/nanopore_CSF_cfDNA

### Competing Interest Statement

The authors have declared no competing interest.

## REFERENCES

1. Nieder C, Haukland E, Mannsaker B, Pawinski AR, Yobuta R, Dalhaug A: Presence of Brain Metastases at Initial Diagnosis of Cancer: Patient Characteristics and Outcome. Cureus 2019, 11:e4113.

2. Schouten LJ, Rutten J, Huveneers HA, Twijnstra A: Incidence of brain metastases in a cohort of patients with carcinoma of the breast, colon, kidney, and lung and melanoma. Cancer 2002, 94:2698–2705.

3. Naresh G, Malik PS, Khurana S, Pushpam D, Sharma V, Yadav M, Jain D, Pathy S: Assessment of Brain Metastasis at Diagnosis in Non-Small-Cell Lung Cancer: A Prospective Observational Study From North India. JCO Glob Oncol 2021, 7:593–601.

4. Kahraman S, Karakaya S, Kaplan MA, Goksu SS, Ozturk A, Isleyen ZS, Hamdard J, Yildirim S, Dogan T, Isik S, et al: Treatment outcomes and prognostic factors in patients with driver mutant non-small cell lung cancer and de novo brain metastases. Sci Rep 2024, 14:5820.

5. Friedlaender A, Perol M, Banna GL, Parikh K, Addeo A: Oncogenic alterations in advanced NSCLC: a molecular super-highway. Biomark Res 2024, 12:24.

6. Shin DY, Na, II, Kim CH, Park S, Baek H, Yang SH: EGFR mutation and brain metastasis in pulmonary adenocarcinomas. J Thorac Oncol 2014, 9:195–199.

7. Bettegowda C, Sausen M, Leary RJ, Kinde I, Wang Y, Agrawal N, Bartlett BR, Wang H, Luber B, Alani RM, et al: Detection of circulating tumor DNA in early- and late-stage human malignancies. Sci Transl Med 2014, 6:224ra224.

8. McEwen AE, Leary SES, Lockwood CM: Beyond the Blood: CSF-Derived cfDNA for Diagnosis and Characterization of CNS Tumors. Front Cell Dev Biol 2020, 8:45.

9. Escudero L, Martinez-Ricarte F, Seoane J: ctDNA-Based Liquid Biopsy of Cerebrospinal Fluid in Brain Cancer. Cancers (Basel) 2021, 13.

10. Deng ZY, Cui L, Li PS, Ren NJ, Zhong Z, Tang Z, Wang L, Gong JW, Cheng HF, Guan YF, et al: Genomic comparison between cerebrospinal fluid and primary tumor revealed the genetic events associated with brain metastasis in lung adenocarcinoma. Cell Death & Disease 2021, 12.

11. Pfeifer GP, Xiong W, Hahn MA, Jin SG: The role of 5-hydroxymethylcytosine in human cancer. Cell Tissue Res 2014, 356:631–641.

12. Li W, Liu M: Distribution of 5-hydroxymethylcytosine in different human tissues. J Nucleic Acids 2011, 2011:870726.

13. Lau BT, Almeda A, Schauer M, McNamara M, Bai X, Meng Q, Partha M, Grimes SM, Lee H, Heestand GM, Ji HP: Single-molecule methylation profiles of cell-free DNA in cancer with nanopore sequencing. Genome Med 2023, 15:33.

14. Liu Y, Rosikiewicz W, Pan Z, Jillette N, Wang P, Taghbalout A, Foox J, Mason C, Carroll M, Cheng A, Li S: DNA methylation-calling tools for Oxford Nanopore sequencing: a survey and human epigenome-wide evaluation. Genome Biol 2021, 22:295.

15. Yuen ZW, Srivastava A, Daniel R, McNevin D, Jack C, Eyras E: Systematic benchmarking of tools for CpG methylation detection from nanopore sequencing. Nat Commun 2021, 12:3438.

16. Li H, Handsaker B, Wysoker A, Fennell T, Ruan J, Homer N, Marth G, Abecasis G, Durbin R, Genome Project Data Processing S: The Sequence Alignment/Map format and SAMtools. Bioinformatics 2009, 25:2078–2079.

17. Frankish A, Diekhans M, Jungreis I, Lagarde J, Loveland JE, Mudge JM, Sisu C, Wright JC, Armstrong J, Barnes I, et al: Gencode 2021. Nucleic Acids Res 2021, 49:D916–D923.

18. de Medeiros Oliveira M, Bonadio I, Lie de Melo A, Mendes Souza G, Durham AM: TSSFinder-fast and accurate ab initio prediction of the core promoter in eukaryotic genomes. Brief Bioinform 2021, 22.

19. Umarov R, Kuwahara H, Li Y, Gao X, Solovyev V: Promoter analysis and prediction in the human genome using sequence-based deep learning models. Bioinformatics 2019, 35:2730–2737.

20. Zhao J, Qian F, Li X, Yu Z, Zhu J, Yu R, Zhao Y, Ding K, Li Y, Yang Y, et al: CanMethdb: a database for genome-wide DNA methylation annotation in cancers. Bioinformatics 2023, 39.

21. Corces MR, Granja JM, Shams S, Louie BH, Seoane JA, Zhou W, Silva TC, Groeneveld C, Wong CK, Cho SW, et al: The chromatin accessibility landscape of primary human cancers. Science 2018, 362.

22. Wang YY, Xu RJ, Zhang DY, Lu T, Yu WP, Wo Y, Liu A, Sui TY, Cui J, Qin Y, et al: Circ-ZKSCAN1 regulates FAM83A expression and inactivates MAPK signaling by targeting miR-330-5p to promote non-small cell lung cancer progression. Translational Lung Cancer Research 2019, 8:862-+.

23. Sun QY, Ding LW, Xiao JF, Chien WW, Lim SL, Hattori N, Goodglick L, Chia D, Mah V, Alavi M, et al: SETDB1 accelerates tumourigenesis by regulating the WNT signalling pathway. Journal of Pathology 2015, 235:559–570.

24. Yang Q, Zhu L, Ye M, Zhang B, Zhan PH, Li H, Zou W, Liu J: Tumor Suppressor 4.1N/ is Epigenetic Silenced by Promoter Methylation and MiR-454-3p in NSCLC. Frontiers in Genetics 2022, 13.

25. Qiao H, Feng Y, Tang HP: COL6A6 inhibits the proliferation and metastasis of non-small cell lung cancer through the JAK signalling pathway. Translational Cancer Research 2021, 10:4514–4522.

26. Xie Z, Bailey A, Kuleshov MV, Clarke DJB, Evangelista JE, Jenkins SL, Lachmann A, Wojciechowicz ML, Kropiwnicki E, Jagodnik KM, et al: Gene Set Knowledge Discovery with Enrichr. Curr Protoc 2021, 1:e90.

27. Liu H, Wu Y, Zhu S, Liang W, Wang Z, Wang Y, Lv T, Yao Y, Yuan D, Song Y: PTP1B promotes cell proliferation and metastasis through activating src and ERK1/2 in non-small cell lung cancer. Cancer Lett 2015, 359:218–225.

28. Hsu PC, Yang CT, Jablons DM, You L: The Crosstalk between Src and Hippo/YAP Signaling Pathways in Non-Small Cell Lung Cancer (NSCLC). Cancers (Basel) 2020, 12.

29. Gao SP, Chang Q, Mao N, Daly LA, Vogel R, Chan T, Liu SH, Bournazou E, Schori E, Zhang H, et al: JAK2 inhibition sensitizes resistant EGFR-mutant lung adenocarcinoma to tyrosine kinase inhibitors. Sci Signal 2016, 9:ra33.

30. Lee SM: Detecting DNA hydroxymethylation: exploring its role in genome regulation. BMB Rep 2024, 57:135-142.

31. He B, Zhang C, Zhang X, Fan Y, Zeng H, Liu J, Meng H, Bai D, Peng J, Zhang Q, et al: Tissue-specific 5-hydroxymethylcytosine landscape of the human genome. Nat Commun 2021, 12:4249.

32. Xue D, Yang P, Wei Q, Li X, Lin L, Lin T: IL-21/IL-21R inhibit tumor growth and invasion in non-small cell lung cancer cells via suppressing Wnt/beta-catenin signaling and PD-L1 expression. Int J Mol Med 2019, 44:1697–1706.

33. Burgess JT, Bolderson E, Adams MN, Duijf PHG, Zhang SD, Gray SG, Wright G, Richard DJ, O’Byrne KJ: SASH1 is a prognostic indicator and potential therapeutic target in non-small cell lung cancer. Scientific Reports 2020, 10.

34. Higgins G, Roper KM, Watson IJ, Blackhall FH, Rom WN, Pass HI, Ainscough JF, Coverley D: Variant Ciz1 is a circulating biomarker for early-stage lung cancer. Proc Natl Acad Sci U S A 2012, 109:E3128–3135.

35. Wang Y, Gong W, Zhou S, Yang L, Qiu F, Lin M, Su W, Nie W, Datta S, Rao B, et al: Long Noncoding RNA PRRG4-4 Promotes Viability, Cell Cycle, Migration, and Invasion in Lung Cancer Cells. DNA Cell Biol 2018, 37:953–966.

36. Pulido I, Ollosi S, Aparisi S, Becker JH, Aliena-Valero A, Benet M, Rodriguez ML, Lopez A, Tamayo-Torres E, Chulia-Peris L, et al: Endothelin-1-Mediated Drug Resistance in EGFR-Mutant Non-Small Cell Lung Carcinoma. Cancer Res 2020, 80:4224–4232.

37. Ding J, Yeh CR, Sun Y, Lin C, Chou J, Ou Z, Chang C, Qi J, Yeh S: Estrogen receptor beta promotes renal cell carcinoma progression via regulating LncRNA HOTAIR- miR-138/200c/204/217 associated CeRNA network. Oncogene 2018, 37:5037–5053.

38. Yeh CT, Chen TT, Satriyo PB, Wang CH, Wu ATH, Chao TY, Lee KY, Hsiao M, Wang LS, Kuo KT: Bruton’s tyrosine kinase (BTK) mediates resistance to EGFR inhibition in non-small-cell lung carcinoma. Oncogenesis 2021, 10:56.

39. Chen H, Moreno-Moral A, Pesce F, Devapragash N, Mancini M, Heng EL, Rotival M, Srivastava PK, Harmston N, Shkura K, et al: WWP2 regulates pathological cardiac fibrosis by modulating SMAD2 signaling. Nat Commun 2019, 10:3616.

40. Wang DG, Gao J, Wang J, Li KC, Wu ZB, Liao ZM, Wu YB: TFAP2A drives non-small cell lung cancer (NSCLC) progression and resistance to targeted therapy by facilitating the ESR2-mediated MAPK pathway. Cell Death Discov 2024, 10:491.

41. Zhao L, Wang Y, Sun X, Zhang X, Simone N, He J: ELK1/MTOR/S6K1 Pathway Contributes to Acquired Resistance to Gefitinib in Non-Small Cell Lung Cancer. Int J Mol Sci 2024, 25.

42. Jiang T, Li N, Xu H, Sun L, Zhang Y, Luo Q, Yang L: Identification of ATAD3A as a key regulator in non-small cell lung cancer by promoting STAT3-induced cell proliferation and tumor angiogenesis. Molecular Carcinogenesis 2024, 63:510–523.

43. Salaun M, Peng J, Hensley HH, Roder N, Flieder DB, Houlle-Crepin S, Abramovici- Roels O, Sabourin JC, Thiberville L, Clapper ML: MMP-13 In-Vivo Molecular Imaging Reveals Early Expression in Lung Adenocarcinoma. PLoS One 2015, 10:e0132960.

44. Yamashita SI, Chujo M, Moroga T, Anami K, Tokuishi K, Miyawaki M, Kawano Y, Takeno S, Yamamoto S, Kawahara K: DYRK2 Expression May be a Predictive Marker for Chemotherapy in Non-small Cell Lung Cancer. Anticancer Research 2009, 29:2753–2757.

45. Li Z, Zhang Y, Jin T, Men J, Lin Z, Qi P, Piao Y, Yan G: NQO1 protein expression predicts poor prognosis of non-small cell lung cancers. BMC Cancer 2015, 15:207.

46. Guo Y, Hou J, Luo Y, Wang D: Functional disruption of macrophage migration inhibitory factor (MIF) suppresses proliferation of human H460 lung cancer cells by caspase-dependent apoptosis. Cancer Cell Int 2013, 13:28.

47. Xu C, Liu D, Mei H, Hu J, Luo M: Knockdown of RAD54B expression reduces cell proliferation and induces apoptosis in lung cancer cells. J Int Med Res 2019, 47:5650–5659.

48. Seki Y, Yoshida Y, Ishimine H, Shinozaki-Ushiku A, Ito Y, Sumitomo K, Nakajima J, Fukayama M, Michiue T, Asashima M, Kurisaki A: Lipase member H is a novel secreted protein selectively upregulated in human lung adenocarcinomas and bronchioloalveolar carcinomas. Biochemical and Biophysical Research Communications 2014, 443:1141–1147.

49. Zhang L, Li JY, Lv XJ, Guo TT, Li W, Zhang J: MID1-PP2A complex functions as new insights in human lung adenocarcinoma. Journal of Cancer Research and Clinical Oncology 2018, 144:855–864.

50. van der Pol Y, Tantyo NA, Evander N, Hentschel AE, Wever BM, Ramaker J, Bootsma S, Fransen MF, Lenos KJ, Vermeulen L, et al: Real-time analysis of the cancer genome and fragmentome from plasma and urine cell-free DNA using nanopore sequencing. EMBO Mol Med 2023, 15:e17282.

51. Qi T, Pan M, Shi H, Wang L, Bai Y, Ge Q: Cell-Free DNA Fragmentomics: The Novel Promising Biomarker. Int J Mol Sci 2023, 24.

52. Cristiano S, Leal A, Phallen J, Fiksel J, Adleff V, Bruhm DC, Jensen SO, Medina JE, Hruban C, White JR, et al: Genome-wide cell-free DNA fragmentation in patients with cancer. Nature 2019, 570:385–389.

53. Mouliere F, Mair R, Chandrananda D, Marass F, Smith CG, Su J, Morris J, Watts C, Brindle KM, Rosenfeld N: Detection of cell-free DNA fragmentation and copy number alterations in cerebrospinal fluid from glioma patients. EMBO Mol Med 2018, 10.

54. Wu X, Xing PY, Shi M, Guo WH, Zhao FP, Zhu HL, Xiao JP, Wan JH, Li JL: Cerebrospinal Fluid Cell-Free DNA-Based Detection of High Level of Genomic Instability Is Associated With Poor Prognosis in NSCLC Patients With Leptomeningeal Metastases. Frontiers in Oncology 2022, 12.

55. White MD, Klein RH, Shaw B, Kim A, Subramanian M, Mora JL, Giobbie-Hurder A, Nagabhushan D, Jain A, Singh M, et al: Detection of Leptomeningeal Disease Using Cell-Free DNA From Cerebrospinal Fluid. JAMA Netw Open 2021, 4:e2120040.

56. Xu YJ, Huang ZY, Yu XQ, Chen KY, Fan Y: Integrated genomic and DNA methylation analysis of patients with advanced non-small cell lung cancer with brain metastases. Molecular Brain 2021, 14.

57. Besaratinia A, Caceres A, Tommasi S: DNA Hydroxymethylation in Smoking-Associated Cancers. Int J Mol Sci 2022, 23.

58. Song CX, Yin SL, Ma L, Wheeler A, Chen Y, Zhang Y, Liu B, Xiong JJ, Zhang WH, Hu JK, et al: 5-Hydroxymethylcytosine signatures in cell-free DNA provide information about tumor types and stages. Cell Research 2017, 27:1231–1242.

59. Liao YF, Gu J, Wu YB, Long X, Ge D, Xu JJ, Ding JY: Low level of 5- Hydroxymethylcytosine predicts poor prognosis in non-small cell lung cancer. Oncology Letters 2016, 11:3753–3760.

60. Chen Y, Cai Q, Pan C, Liu W, Li L, Liu J, Gao M, Li X, Wang L, Rao Y, et al: CDK2 Inhibition Enhances Antitumor Immunity by Increasing IFN Response to Endogenous Retroviruses. Cancer Immunol Res 2022, 10:525–539.

61. Peng J, Yang S, Ng CSH, Chen GG: The role of FOXP3 in non-small cell lung cancer and its therapeutic potentials. Pharmacol Ther 2023, 241:108333.

62. Remsik J, Boire A: The path to leptomeningeal metastasis. Nat Rev Cancer 2024, 24:448–460.

63. Guler GD, Ning YH, Coruh C, Mognol GP, Phillips T, Nabiyouni M, Hazen K, Scott A, Volkmuth W, Levy S: Plasma cell-free DNA hydroxymethylation profiling reveals anti-PD-1 treatment response and resistance biology in non-small cell lung cancer. Journal for Immunotherapy of Cancer 2024, 12.

64. Maansson CT, Helstrup S, Ebert EBF, Meldgaard P, Sorensen BS: Circulating immune response proteins predict the outcome following disease progression of osimertinib treated epidermal growth factor receptor-positive non-small-cell lung cancer patients. Translational Lung Cancer Research 2023, 12:14-+.

65. Shiiya A, Noguchi T, Tomaru U, Ariga S, Takashima Y, Ohhara Y, Taguchi J, Takeuchi S, Shimizu Y, Kinoshita I, et al: EGFR inhibition in EGFR-mutant lung cancer cells perturbs innate immune signaling pathways in the tumor microenvironment. Cancer Science 2023, 114:1270–1283.

66. Shang YH, Li XF, Liu WW, Shi XL, Yuan SH, Huo R, Fang GT, Han X, Zhang JN, Wang KJ, et al: Comprehensive genomic profile of Chinese lung cancer patients and mutation characteristics of individuals resistant to icotinib/gefitinib. Scientific Reports 2020, 10.

