## Supplementary information for "Nanopore-based cell-free DNA fragmentation and methylation profiles from the cerebral spinal fluid of patients with lung cancer brain metastases"

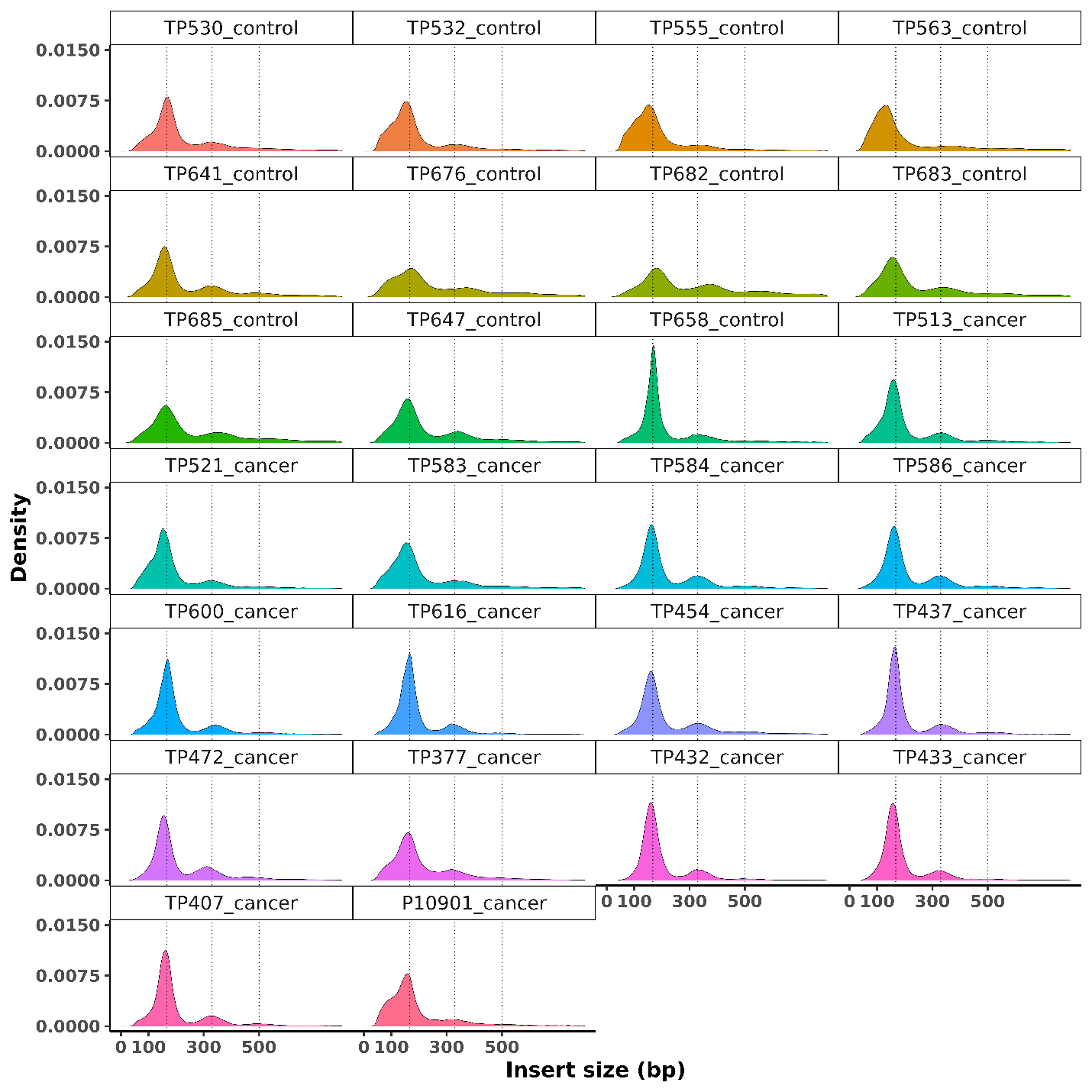

**Supplementary Figure 1. cfDNA fragment size distribution in CSF from healthy donors and NSCLC-BMET cancer patients.** The cfDNA fragment size distribution of healthy control CSF (N=11, labeled as ‘control’) and cancer patient-derived CSF (N=15, labeled as ‘cancer’) were shown. Dotted lines indicated mono- (167 bp), di- (330 bp) and tri-nucleosomes (500 bp).

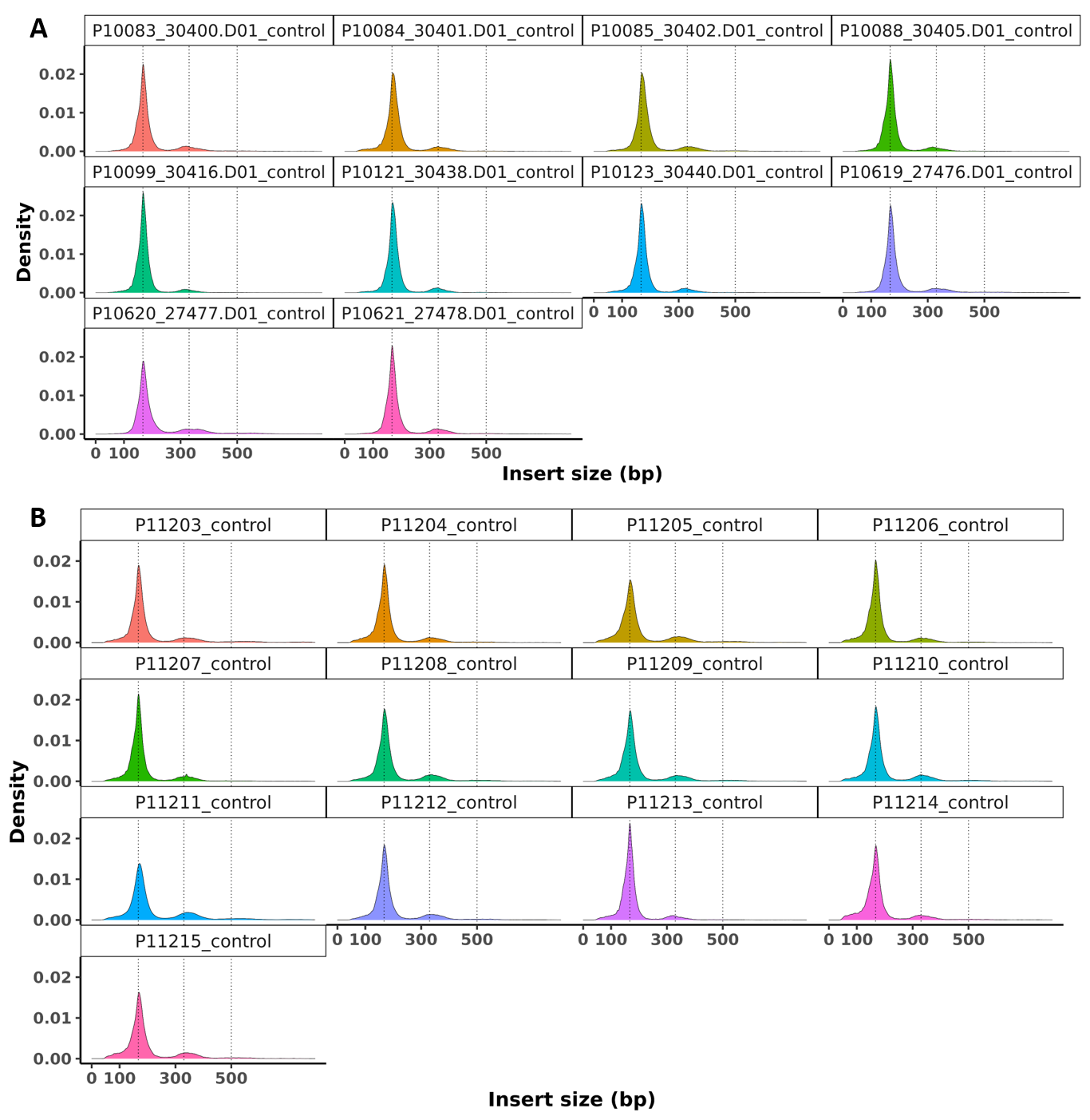

**Supplementary Figure 2. cfDNA fragment size distribution in plasma from two independent sources of healthy donors.** A) The cfDNA fragment size distribution of healthy control plasma (N=10) from Precision For Medicine was shown. 10 samples were randomly selected out of 198 to show the distribution. B) The cfDNA fragment size distribution of healthy control plasma (N=13) from Discovery Life Sciences was shown. Dotted lines indicated mono- (167 bp), di- (330 bp) and tri-nucleosomes (500 bp).

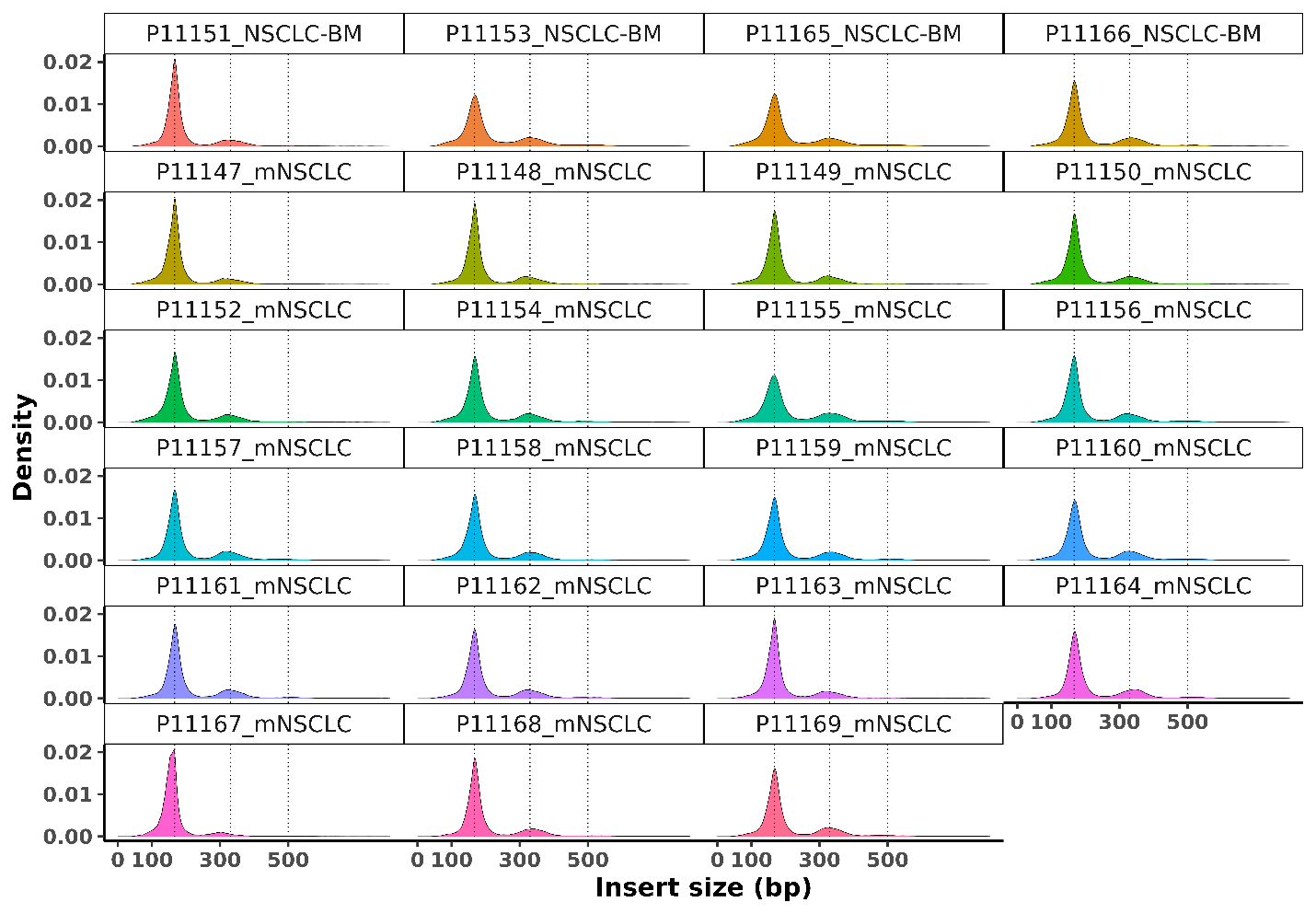

**Supplementary Figure 3. cfDNA fragment size distribution in plasma from metastatic NSCLC cancer patients.** The cfDNA fragment size distribution of metastatic NSCLC patient-derived plasma (N=23) from Discovery Life Sciences was shown. The cohort included 4 NSCLC-BMET patients (labeled as ‘NSCLC-BM’) and 19 metastatic NSCLC patients with other sites (labeled as ‘mNSCLC’). Dotted lines indicated mono- (167 bp), di- (330 bp) and tri-nucleosomes (500 bp).

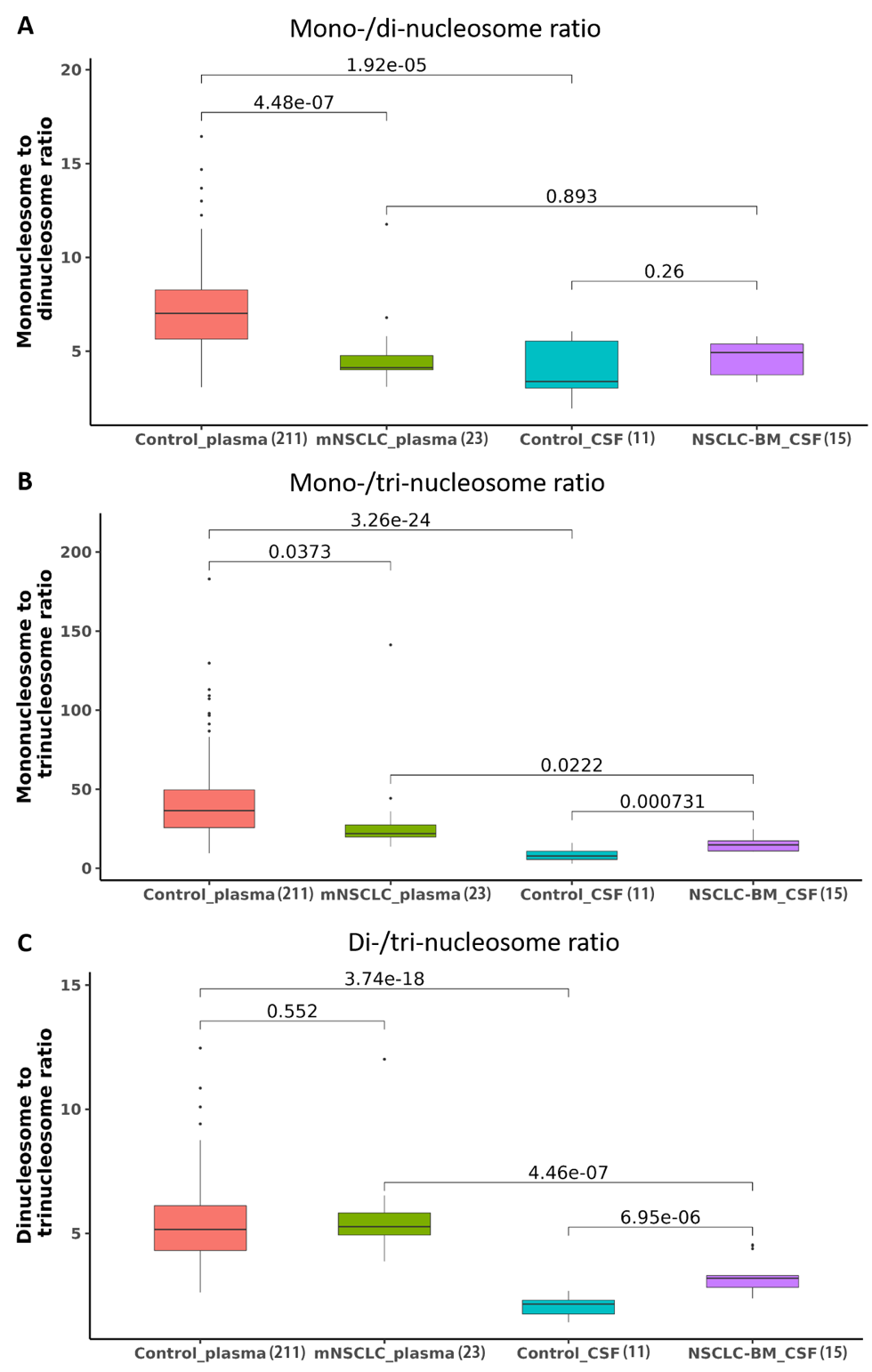

**Supplementary Figure 4. cfDNA nucleosome ratios in CSF and plasma from NSCLC-BMET patients and healthy controls.** A) The mono-/di-nucleosome ratios of healthy control plasma (N=211, 2 vendor sources combined), metastatic NSCLC patient-derived plasma (N=23), healthy control CSF (N=11), and NSCLC-BMET patient-derived CSF (N=15). B) The mono-/tri-nucleosome ratios of healthy control plasma (N=211, 2 vendor sources combined), metastatic NSCLC patient-derived plasma (N=23), healthy control CSF (N=11), and NSCLC-BMET patient-derived CSF (N=15). C) The di-/tri-nucleosome ratios of healthy control plasma (N=211, 2 vendor sources combined), metastatic NSCLC patient-derived plasma (N=23), healthy control CSF (N=11), and NSCLC-BMET patient-derived CSF (N=15).

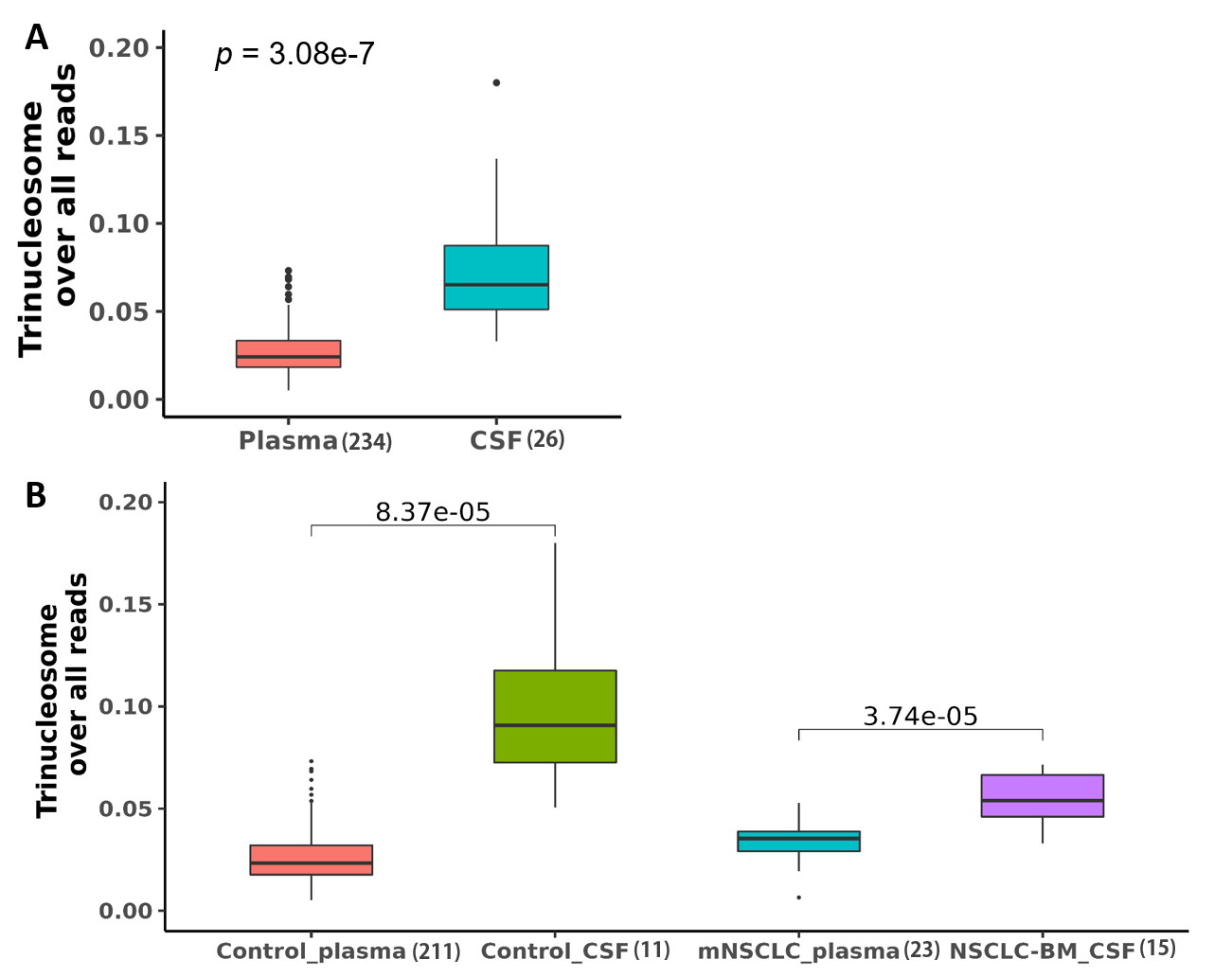

**Supplementary Figure 5. cfDNA trinucleosome fraction in CSF and plasma from NSCLC-BMET patients and healthy controls.** The normalized tri-nucleosome fraction over combination of mono-, di-, and tri-nucleosomes in A) plasma (N=234) vs. CSF (N=26) overall, and B) healthy control plasma from healthy control plasma (N=211, 2 vendor sources combined), healthy control CSF (N=11), metastatic NSCLC patient-derived plasma (N=23), and NSCLC-BMET patient-derived CSF (N=15).

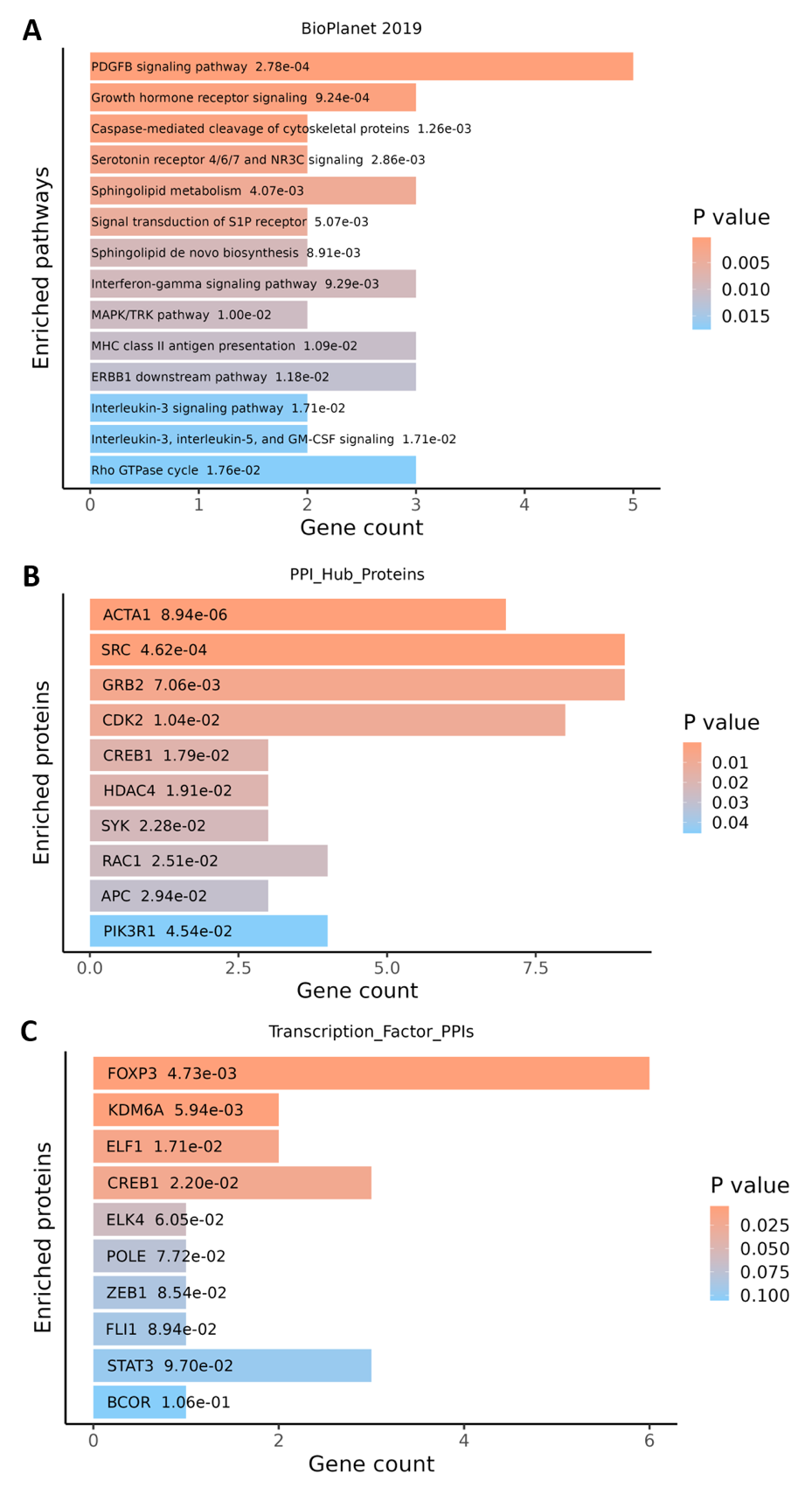

**Supplementary Figure 6. Analysis of enriched pathways and protein-protein interactions for promoter 5hmCG in cancer patients.** A) Pathway enrichment analysis was performed by Enrichr using BioPlanet 2019. p-values were shown for each associated pathway. B) Protein-protein interaction analysis was performed by Enrichr using PPI Hub Proteins database. C) Protein-protein interaction analysis was performed by Enrichr using Transcription Factor PPIs database. p-values were shown for each associated protein.
